# transfactor: Transcription factor activity estimation via probabilistic gene expression deconvolution

**DOI:** 10.1101/2025.03.19.644088

**Authors:** Koen Van den Berge, Peter Bickel, Sandrine Dudoit

**Affiliations:** Department of Statistics, University of California, Berkeley, CA, USA; Department of Applied Mathematics, Computer Science and Statistics, Ghent University, Belgium; Graduate Group in Biostatistics, University of California, Berkeley, CA, USA; Division of Biostatistics, School of Public Health, University of California, Berkeley, CA, USA; Center for Computational Biology, University of California, Berkeley, CA, USA

## Abstract

Gene expression is the primary modality being studied to differentiate between biological cells. Contemporary single-cell studies simultaneously measure genome-wide transcription levels for thousands of individual cells in a single experiment. While the characterization of cell population differences has often occurred through differential expression analysis, tiny effect sizes become statistically significant when thousands of cells are available for each population, compromising biological interpretation. Moreover, these large studies have spurred the development of methods to infer gene regulatory networks (GRNs) directly from the data, and GRN databases are becoming more comprehensive.

In this work, we propose a statistical model for gene expression measures and an inference method that leverage GRNs to deconvolve transcription factor (TF) activity from gene expression, by probabilistically assigning mRNA molecules to TFs. This shifts the paradigm from investigating gene expression differences to regulatory differences at the level of TF activity, aiding interpretation and allowing prioritization of a limited number of TFs responsible for significant contributions to the observed gene expression differences. The inferred TF activities result in intuitive prioritization of TFs in terms of the (difference in) estimated number of molecules they produce, in contrast to other widely-used methods relying on arbitrary enrichment scores. Our model allows the incorporation of prior information on the regulatory potential between each TF and target gene through prior distributions, and is able to deal with both repressing and activating interactions. We compare our approach to other TF activity estimation methods using two simulation experiments and two case studies.

## 1 Introduction

In single-cell RNA sequencing (scRNA-seq), researchers investigate gene expression levels as driving factors of observed biological differences between cell types, disease states, and responses to treatments. The resolution of gene expression at the level of single cells has been a boon for biological research, as it has allowed to deconvolve cellular gene expression heterogeneity, as well as answer a new set of questions, including the identification of homogeneous cell entities (‘cell types’) or dynamic cellular differentiation processes (‘trajectories’) such as cell development and activation.

Traditionally, differences between cell types or within/between trajectories have been identified using differential expression (DE) analysis, where gene-level differences in average expression are assessed using hypothesis testing [Finak et al., 2015, Van den Berge et al., 2020]. Through this approach, the study and interpretation of differentially expressed genes (DEG) therefore provides an opening to elucidate the biological processes of interest. However, the large number of sequenced cells (in the tens or hundreds of thousands) of contemporary scRNA-seq datasets renders biologically irrelevant (or insignificant) effect sizes statistically significant. While moderation methods such as those implemented in the `limma` framework [Smyth, 2004] were designed in part to help with the issue of biologically uninteresting statistical significance, they are unable to fully resolve the problem. This hampers interpretation and leads to ad hoc tuning of significance thresholds as low as 10^*—*12^ [McFaline-Figueroa et al., 2019]. Even then, the fraction of genes that are in the set of DEG is often too large for reasonable interpretation by biologists. One possible solution is to test against a fold-change threshold [McCarthy and Smyth, 2009a], but this again relies on the ad hoc selection of an appropriate threshold for each hypothesis being assessed.

Gene expression is a molecular process that is being regulated by transcription factors (TFs), that is, proteins that bind DNA regions to be transcribed, resulting in the production of pre-messenger RNA (pre-mRNA) molecules, in the case of an activating regulator, or the inhibition of the production (e.g., through spheric hindrance) of premRNA molecules, in the case of a repressing regulator. A more parsimonious way to interpret gene expression differences is to move one level up in the hierarchy: instead of identifying which genes are differentially expressed, identify transcription factors that are driving the observed gene expression differences. Since there are far fewer TFs than genes, this leads to a more manageable interpretation in the hands of biologists. For example, a recent in-depth review by [Lambert et al.2018] provides a manually-curated list of human TFs and identified 1,639 proteins to be “known or likely TFs”, while the current human genome annotation corresponds to around 20,000 protein-coding genes [Frankish et al., 2019]. However, while TF protein abundance is typically high in single cells, the RNA concentration of the respective TF genes is often low [Weinreb et al., 2020, Chen et al., 2021], and the activity of TF proteins may be modulated through biochemical processes such as phosphorylation [?]. It is thus possible that TFs that are highly active, i.e., producing many RNA molecules from their downstream target genes, have relatively low RNA concentration. This effectively jeopardizes direct TF activity estimation using the corresponding TF’s RNA transcripts, motivating statistical methods to estimate TF activity using downstream target genes instead.

Several methods have previously used (sc)RNA-seq data for identifying TFs responsible for driving gene expression differences, while others focus on alternative data modalities such as single-cell chromatin conformation data [Berest et al., 2019]. Gene-expression-based methods typically either first estimate a gene regulatory network (GRN) or require one as input. A GRN specifies the genes being (positively or negatively) regulated by each TF. A set of genes being regulated by one TF is also often called a *regulon*. Some methods, such as TFEA [Rubin et al., 2020], start from a list of regions of interest based on, for example, chromatin conformation or ChIP-seq data, and perform an enrichment analysis to assess which motifs are more abundant in the regions of interest than what could be expected by chance. Other methods, such as `viper` [Alvarez et al., 2016] and `AUCell` [Aibar et al., 2017], use a gene regulatory network to estimate TF activity for each TF directly using the gene expression data. For all three methods, however, the returned activity scores are on arbitrary scales and are therefore not directly interpretable.

In this manuscript, we introduce a new method that uses scRNA-seq data to infer TF activity in terms of readily interpretable estimates of the number of mRNA molecules produced by each TF for each gene in a particular cell. The method, called transfactor, requires a matrix of gene expression measures and a gene regulatory network as input, and probabilistically deconvolves overall gene expression measures into TF-specific gene expression measures that reflect the allocation of transcripts to TFs that may have produced them. transfactor is capable of handling both activating as well as repressing edges in a GRN. A Bayesian extension of the model allows to incorporate prior knowledge on the confidence of each edge in the network, while also allowing the estimation of these prior parameters through an empirical Bayes framework. If many edges are expected to be irrelevant for the dataset being studied (for example, when a GRN from a global database is used, not all edges may be relevant for the current experiment), a sparse initialization step of the fitting algorithm ensures sparse results in terms of TF activity measures, effectively boosting TF activity estimation accuracy in these scenarios. We compare several variants of transfactor to state-of-the-art methods, as identified in Holland et al. [2020], using two simulation experiments, and provide two extensive case study analyses showing how TF activity estimation can unlock biological insight.

## 2. Methods

### 2.1 Statistical model

#### 2.1.1 Notation and definitions

We assume that we know a gene regulatory network that is relevant for the system under study, which comprises *G* genes and *T* transcription factors. We define the *G × T* adjacency matrix A of the GRN, with elements *A*_*gt*_ = 1 (*A*_*gt*_ = —1) if gene *g* is positively (negatively) regulated by TF *t* and zero otherwise. In Section 2.2.2, we show how to account for repressions (negative *A*_*gt*_) prior to model fitting; in the remainder of the manuscript, we will assume there are no repressing edges in A, i.e., only non-negative entries. Based on A, we know that (i) a gene *g* is regulated by a known set of TFs which is denoted by 𝒯_*g*_ = {*t* : *A*_*gt*_ = 1}, and (ii) a TF *t* regulates a known set of genes that is denoted by 𝒢_*t*_ = {*g* : *A*_*gt*_ = 1}.

We assume that cells have been grouped into relevant entities, be it cell classes, states, or experimental conditions, generally corresponding to their cell identity, which we refer to using the general moniker of ‘cell type’. Let *n* denote the total number of observed cells and *n*_*c*_ the number of cells observed for cell type *c, c* = 1,…, *C*. For a particular cell *i* that belongs to cell type *c*(*i*), let *Z*_*gti*_ denote a measure of the number of transcripts of gene *g* produced by TF *t* and let 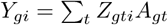 denote a measure of the total number of transcripts of gene *g*. While *Y*_*gi*_ is observed in scRNA-seq, *Z*_*gti*_ is not, and the goal of our method is to deconvolve *Z*_*gti*_ from *Y*_*gi*_. Let *µ*_*gtc*_ and *µ*_*g*.*c*_ denote, respectively, the expected number of transcripts of gene *g* produced by TF *t* in cell type *c* and the expected total number of transcripts of gene *g* in cell type *c*. Note that

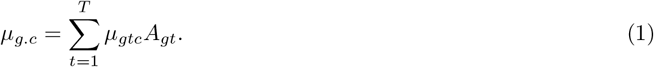

The parameter of interest is *π*_*gtc*_, the proportion of transcripts of gene *g* produced by TF *t* in cell type *c*, i.e.,

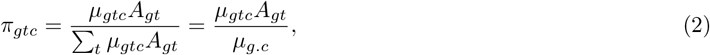

where 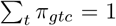 for each gene *g* and cell type *c*. We further introduce 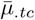 as the global (i.e., across all *g* ∈ 𝒢_*t*_) activity of a TF *t* in cell type *c*, defined as the average number of molecules the TF produces across all the genes it is regulating, i.e.,

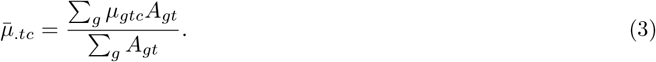

##### Distance-based ranking of transcription factors

In order to identify which TFs may contribute most to differences in expression between cell types, we can construct a distance-based ranking of transcription factors based on their activity. Here, we focus on the Euclidean distance, but note that other distance measures could similarly be used.

A measure of the contribution of a particular TF *t* to differences in gene expression between cell types *c* and *d* can be defined as

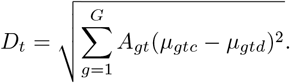

To adjust for gene expression magnitude, we can also work with a scaled distance measure

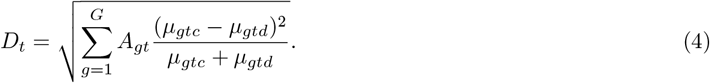

Either measure can then be used to rank TFs according to their contribution to differences in expression between cell types. If a comparison across multiple groups is of interest (e.g., in the olfactory epithelium case study), one could sum *D*_*t*_ across all relevant comparisons or look at contrasts of the *µ*_*gtc*_.

Note that *D*_*t*_ is a more informative measure of the contribution of a particular TF *t* as compared to the Euclidean distance on 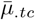 directly. Indeed, *D*_*t*_ also accounts for cases where 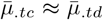, but where *π*_*gtc*_ ≠ *π*_*gtd*_, i.e., the TF is similarly active overall across cell types, but it is preferentially regulating different sets of genes in the different cell types.

#### 2.1.2 Basic model assumptions

For the model as introduced in Section 2.1.1, the parameters *µ*_*gtc*_ are unidentifiable based on only the observed data *Y*_*gi*_. Here, we make two assumptions that should aid towards identifiability. First, we assume a multiplicative model for *µ*_*gtc*_, parameterized according to a gene-specific effect *α*_*gc*_ and a TF-specific effect *β*_*tc*_, i.e.,

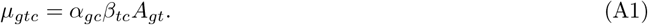

Biologically, (A1) means that, for a given gene *g*, we make the strong assumption that all TFs *t* ∈ 𝒯_*g*_ have the same binding capacity and therefore the competition between different TFs to produce mRNA molecules is only affected by *β*_*tc*_. This is a simplification of reality, where other determinants, such as chromatin conformation, histone marks, and the presence of other TFs, would also affect the TF-specific contribution at a genomic locus [Bannister and Kouzarides, 2011, Sönmezer et al., 2021].

Further, we assume that

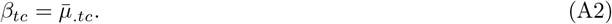

This assumption also helps in making the model parameters interpretable, since we can consider the TF-specific contribution in (A1) to correspond to the global activity of that TF.

#### 2.1.3 Implications of model assumptions

Our model assumptions, (A1) and (A2), have the following implications that are useful to note explicitly, since they aid in interpreting the model as well as understanding its constraints.

First, we note that Equations (3) and (A1) imply that

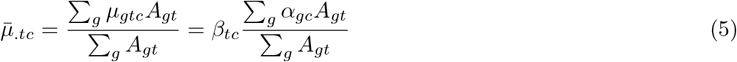

and thus, given (A2), the following constraint must hold

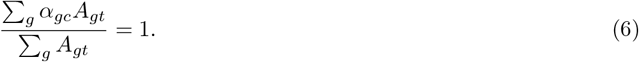

From Equations (1), (A1), and (A2), it follows that

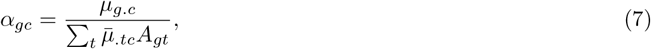

which allows us to rewrite (A1) in a more readily interpretable and intuitive form

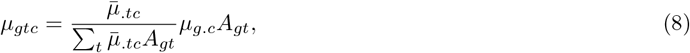

which shows that the contribution of a TF *t* to the gene expression of gene *g* is related to its activity in that cell type 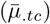 as compared to the activity of all other TFs regulating the gene 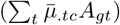. This also allows us to relate *π*_*gtc*_ to the TF activity 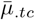, as Equations (2) and (8) imply that

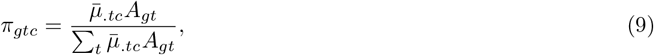

i.e., the proportion of transcripts of gene *g* produced by TF *t* is only dependent on the TF’s global activity 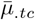, relative to the activity of all other TFs regulating the gene.

2.1.4 Model identifiability

We have not established model identifiability and the proof is a non-trivial mathematical question that is beyond the scope of this paper. The identifiability problem is shared by other methods that analogously attempt to assign k-mers of sequencing reads to transcripts [2016, 2017] (instead of mRNA molecules to TFs). In practice, using small network examples (Supplementary Methods Section 6), we found a few conditions that are necessary, but not sufficient, for model identifiability:

1. *T* ≤ *G*. The number of transcription factors must not exceed the number of genes. This condition should always be met, as it is a biological reality.
2. A must be of full column rank, i.e., column rank *T*. This condition may or may not hold, depending on the provided GRN. If a limited set of TFs are linearly related in terms of their regulated genes (i.e., the corresponding columns in the adjacency matrix are not linearly independent), one may consider removing these TFs in order to obtain a full-column-rank matrix.

#### 2.1.5 Poisson model and a Bayesian extension of the model

The basic assumptions (A1) and (A2) of Section 2.1 naturally lead to a method-of-moments estimation approach, with estimating equations

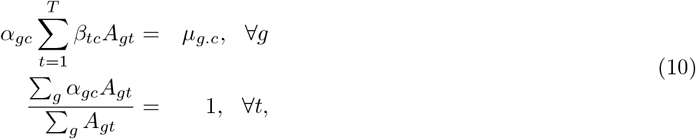

which have no closed-form solution for the the parameters *α*_*gc*_ and *β*_*tc*_. We will not directly use method-of-moments estimation and will instead introduce additional parametric assumptions that allow us to develop a Bayesian extension of the model that incorporates prior biological knowledge and induces sparsity in the parameter estimates.

We assume that the latent variables follow a Poisson distribution, i.e.,

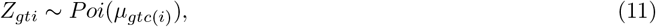

where the *Z*_*gt*i_ are independently distributed across genes, TFs, and cells. This frequentist model will be referred to as the ‘Poisson model’. Note, however, that it does not allow incorporating a priori known information on the edges present in the GRN. Indeed, different edges might have different confidence scores based on empirical evidence supporting each edge, e.g., as obtained via databases or experiments. To incorporate such information, we specify, for each gene *g* and cell type *c*, a Dirichlet prior distribution for (*π*_*gtc*_ : *t* = 1,…, *T*), subject to the constraint of Equation (9), i.e.,

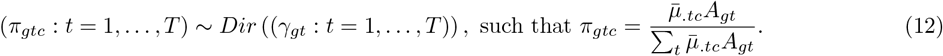

The (*π*_*gtc*_ : *t* = 1,…, *T*) are assumed to be independently distributed over genes *g* and cell types *c* and the prior distributions are constant across cell types. Further, if *A*_*gt*_ = 0, then γ_*gt*_ = 0. This formulation also implies that if genes *g*_1_ and *g*_2_ are regulated by the same TF *t*, then 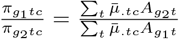, and if they are regulated by exactly the same set of TFs ∀*t*), then 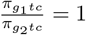.

The constraint on the Dirichlet distribution that the *π*_*gtc*_ have this specific form makes it explicit that there is no inherent sampling variability associated with the Dirichlet prior, but that it is mainly there to borrow information across cell types and/or to incorporate prior knowledge.

In our model, the γ_*gt*_ can be assumed to be either: (i) equal to zero, corresponding to a non-informative (improper) prior distribution, which reduces to inference based on the frequentist Poisson model; (ii) known (e.g., by using information from biological databases on the regulatory strength between a gene and a TF), in which case we are working with a Bayesian model, or (iii) estimated from the data in an empirical Bayes fashion.

The model is thus

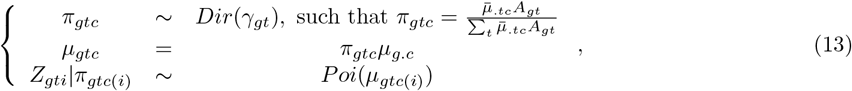

where it follows that *Y*_*gi*_ ∼ *Poi*(*µ*_*g*.*c*(*i*)_) (due to independence over *t*) and also 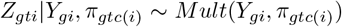. The parameters of interest are *π*_*gtc*_. Note that while downstream interpretation may be focused on either *π*_*gtc*_ or 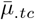, both are readily available once we have estimated one or the other, see Equation (9). We will also estimate γ_*gt*_, if they are not assumed to be known.

### 2.2 EM algorithm for fitting the hierarchical Poisson model

We first note that the parameter *µ*_*g*.*c*_ can be inferred solely from the observed data **Y** = {*Y*_*gi*_ : *i* = 1,…, *n*; *g* = 1,…, *G}* and trivially estimated using the sufficient statistic 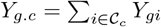, with *C*_*c*_ = {*i* : *c*(*i*) = *c}*, by

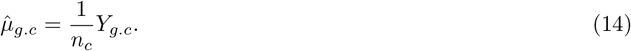

We then use the expectation-maximization (EM) algorithm [Dempster et al., 1977] to estimate the parameters *θ* = {*π*_*gtc*_, *γ*_*gt*_}. In the frequentist setting, the algorithm is used to maximize the likelihood function with respect to *θ* (which, in this case, does not include *γ*), and in the (empirical) Bayes setting, it is used to maximize the posterior density with respect to *θ* (which possibly includes *γ* as unknown).

The frequentist EM algorithm maximizes the *Q*-function

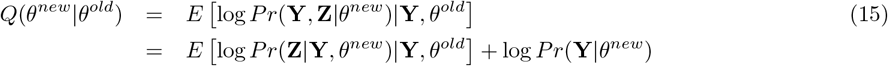

with respect to *θ*^*new*^, where **Z** = {*Z*_*gti*_ : *i* = 1,…, *n*; *g* = 1,…, *G*; *t* = 1,…, *T}*. Note that the second term only involves the parameters *µ*_*g*.*c*_, that have already been inferred from the observed data. Hence, the EM algorithm is only concerned with the first term.

In a Bayesian setting, the EM algorithm can be adapted to estimate the mode of the posterior distribution of *θ* by updating the Q-function in the M-step.

Specifically, the EM algorithm proceeds as follows.

- **E-step:** Calculate *Q* by deriving the expected value of *Z*_*gti*_, given the data Y and current estimates *θ*^*old*^.
- **M-step:** In a Bayesian setting, maximize *Q*(*θ*^*new*^|*θ*^*old*^)+*G*(*θ*^*new*^) with respect to *θ*^*new*^, where *G* is the log-prior density [Dempster et al., 1977]. In our case, *G* corresponds to the log-prior density of the *π*_*gtc*_ parameters, which is assumed to be a Dirichlet density with parameters *γ*_*gt*_.
  1. **Known prior**. Maximize the function *Q*(*θ*^*new*^|*θ*^*old*^) + *G*(*θ*^*new*^) with respect to *π*_*gtc*_, keeping *γ*_*gt*_ fixed. This corresponds to calculating 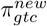.
  2. **Unknown prior**. In an empirical Bayes setting, where we also estimate the parameters of the prior distribution, we additionally maximize the function *Q*(*θ*^*new*^|*θ*^*old*^)+ *G*(*θ*^*new*^) with respect to *γ*_*gt*_, keeping *π*_*gtc*_ fixed at its new value. There is no closed form solution for this, but we will use an approximate MLE as update step. This corresponds to calculating 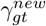.

After each iteration, we set *θ*^*old*^ = *θ*^*new*^. Full details on the EM algorithm and update steps are provided in Supplementary Methods.

Note that, if the prior parameters *γ*_*gt*_ are assumed known, then setting all *γ*_*gt*_ = 1 is equivalent to providing no prior information, see Equation (20) in Supplementary Methods.

In general, it is not guaranteed that the EM algorithm will converge to the global maximum of the likelihood function, and it is possible that the algorithm will instead converge to a local maximum close to its initialization value. However, initializing the EM algorithm with different random starting values in our simulation studies leads to the same parameter estimates upon convergence, providing empirical evidence of good behavior. We recommend that users investigate diagnostic plots to check convergence of the EM algorithm and, in particular, track the log-likelihood evolution across iterations. Our software package transfactor provides functionality for assessing convergence of the EM algorithm (Section 2.3).

#### 2.2.1 Initialization of the EM algorithm

The default initialization for *π*_*gtc*_ is 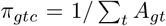, i.e., an equal partitioning of a gene’s transcripts among its regulating TFs. However, while a GRN may indicate that a gene is possibly regulated by many TFs, it is unlikely that all of these plausible TFs are actively contributing to gene expression in a specific biological tissue or condition. It would therefore make sense to obtain sparse estimates of TF activity, where some TFs do not contribute at all to the expression of some genes. If there is prior information, one may promote such sparsity through the Dirichlet prior; indeed, letting *γ*_*gt*_ = 0 will enforce TF *t* to not contribute to the gene expression of gene *g*. However, this is enforced across all cell types, which could be suboptimal. In addition, prior information to promote such sparsity may not be available. An alternative is to promote sparsity in the TF activity estimates in an unsupervised manner. There are several approaches to do so. A natural paradigm would be to add a sparsity-inducing penalty to the log-likelihood, to be incorporated in the fitting using the EM algorithm. However, many of these approaches result in non-closed-form solutions in the M-step, often requiring numerical optimization at each iteration of the EM algorithm, thereby drastically increasing the computational burden of model fitting.

Instead, we use a sparse initialization of the parameters in the EM algorithm. Indeed, if a particular 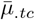 is initialized as 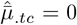, then the update steps of the EM algorithm will keep it zero provided *γ*_*gt*_ = 1 (i.e., if we do not feed prior information to the EM algorithm through g, see Equation (20) in Supplementary Methods) or if we use the Poisson model without the Dirichlet prior. In order to derive these sparse initial estimates, we rely on a Gaussian approximation to the Poisson model, as detailed below.

##### A Gaussian approximation to the Poisson model to promote unsupervised sparse solutions

The sufficient statistics for *µ*_g.*c*_ are *Y*_g.*c*_. We rely on a Gaussian approximation to the Poisson model, where

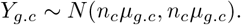

Note that, given Equation (9) and 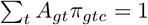,

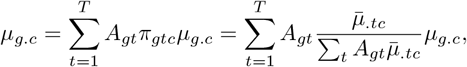

where, as before, we are interested in estimating 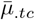 (and, correspondingly, *π*_*gtc*_).

The minimum modified Chi-squared objective function to estimate 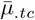, according to Neyman [1949], corresponds to

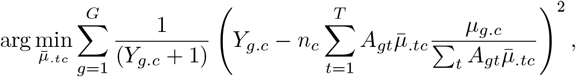

where we use a heuristic pseudocount of 1 in the denominator to avoid division by zero. In order to provide sparse initial estimates, we further use a LASSO approach [Tibshirani, 1996], minimizing the penalized empirical risk for each cell type separately,

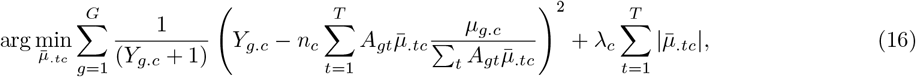

subject to 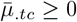. In the loss function, *λ*_*c*_ is a penalty parameter that controls the amount of shrinkage and that is estimated using cross-validation for each cell type separately. The R package glmnet is used for fitting the regression model using the LASSO [Friedman et al., 2010]. The solutions for each cell type will be our initial estimates 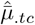 which will be provided to the EM algorithm.

###### Remark 1

Note that the solutions 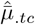 will be sparse, however, differently so for each cell type, meaning that some TFs will contribute to gene expression in some cell types but not in others.

###### Remark 2

If no prior information is provided, the LASSO initialization above will be decisive in terms of which TFs do not contribute to gene expression within a particular cell type. Indeed, if 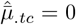, then 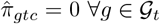.

###### Remark 3

If prior information is provided, i.e., if *γ*_*gt*_ *>* 1, then the sparsity of the LASSO will be ‘overridden’ by the prior information, see Equation (20) in Supplementary Methods.

#### 2.2.2 Accounting for repressions

Our method attempts to assign mRNA molecules to the TFs that produced them. If the GRN contains repressing edges between a TF and a target gene that itself is not a TF, then we discard these interactions (i.e., set *A*_*gt*_ = 0). These repressing edges may have hampered the production of molecules for the specific target gene, but this is not something that we attempt to model. Instead, we are only interested in deconvolving counts for molecules that have actually been produced. In contrast, we do account for repressing interactions between TFs, as these may be informative to assign produced molecules to them. Indeed, if a TF’s repressor is highly active, it is less likely that this particular TF will have produced molecules (i.e., it will have a lower TF activity 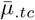).

In order to account for this type of repression, we apply the following procedure prior to fitting the (hierarchical) Poisson model. A TF that is itself being repressed by other TFs will occur both as a row and as a column in the adjacency matrix **A**. We denote the row of **A** corresponding to the gene *g*_*t*_ for TF *t* by 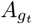 and the set of TFs that are repressing TF *t* by 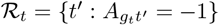. Also, let 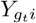 denote the (observed) gene expression measure of repressed TF *t* in cell *i* from cell type *c*(*i*) and 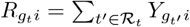 denote the sum of the gene expression measures across all TFs that are repressing TF *t*. If, in the system under study, TF *t* is actually repressed by (a subset of) the TFs in *R*_*t*_, we expect 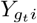 to be negatively associated with 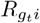. We model 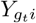 as a function of log 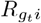 using a Poisson generalized linear model

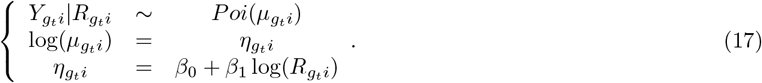

In this model, a negative *β*_1_ corresponds to repressing interactions. We choose to account for repressing interactions if *β*_1_ is found to be significantly different from zero at nominal significance level of 5% after fitting the model in Equation (17).

Repressing interactions are then handled by multiplying 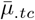 using a scaling factor *ρ*_*tc*_ ∈ [0, 1] estimated as

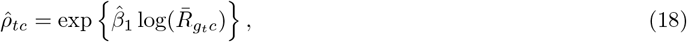

reflecting the ratio of the expression of TF *t* given the average expression of its repressing TFs relative to no expression of its repressing TFs. Here,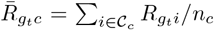. See Supplementary Methods for details on how we use *ρ*_*tc*_ in the model fitting.

### 2.3 Software implementation

The proposed methodology is implemented in the open-source R software package transfactor, which will be submitted to the Bioconductor Project. TF activity can be estimated using the (hierarchical) Poisson model with or without information for the Dirichlet prior distribution. In the fitting of the EM algorithm, we work with sufficient statistics *Y*_g.*c*_ to ease the burden on computational memory requirements.

### 2.4 Benchmarking and case studies

#### 2.4.1 Evaluated methods

We evaluate three different variants of our method:

1. `poisson:` Poisson model.
2. `dirMult:` Hierarchical Poisson model with known Dirichlet prior.
3. `dirMultEBayes:` Hierarchical Poisson model with unknown Dirichlet prior.

For each of these variants, the EM algorithm is initialized either with or without the sparse LASSO-based approach described in Section 2.2.1, which we indicate by the suffix `lasso`, leading to a total of six different methods to evaluate. Except for the Gamma-model-based simulation framework from Section 2.4.3, where no repressing interactions are present, we also evaluate the effect of accounting for repressions as described in Section 2.2.2, which we indicate by the suffix `repr`.

As competing methods, we also evaluate `viper` [Alvarez et al., 2016] and `AUCell` [Aibar et al., 2017], which are, top-performing methods according to the benchmarking study of and Holland et al. [2020]. `viper`. We use `viper` v1.24.0. We run a single-sample analysis, where TF activity is estimated for each sample (cell) independently, by calculating normalized enrichment scores using the `scale` method. In brief, `viper` compares the gene expression of each sample to the average across all samples. The expression counts for each gene are standardized to have zero mean and unit standard deviation across cells. A rank-based enrichment score is computed, where it is assumed that a TF is active if its targets have high ranks (i.e., are highly expressed). The algorithm is capable of incorporating both activating as well as repressing edges, and also information on the confidence of each edge between a TF and a target gene. We run `viper` using the same confidence scores as the dirMult model of transfactor, effectively giving it an advantage as compared to other methods not using these confidence scores.

`AUCell`. We use `AUCell` v1.12.0. Like `viper`, `AUCell` also first scales the expression counts to a gene-wise mean of zero and standard deviation of one across all cells/samples before calculating TF activity scores. `AUCell` calculates TF activity scores by ranking the (by default, top 5%) scaled gene expression measures within each cell, and calculating the area under the receiver operating characteristic (ROC) curve for each regulon (i.e., set of genes regulated by a TF), where a ‘true positive’ is considered a gene that belongs to the regulon. The enrichment score thus reflects the presence of genes from the regulon in the top-expressed genes in the cell.

The main difference between `transfactor` and competing methods `AUCell` and `viper`, is that `transfactor` is based on a statistical model that aligns with the data-generating mechanism and makes clear what the assumptions are. Through this, the parameter estimates of `transfactor` are interpretable; for each gene, one can investigate the relative contribution of TFs to its expression, and how this possibly varies across conditions.

#### 2.4.2 Differential activity analysis

In order to identify TFs that are differentially active under different conditions, we first obtain the estimated number of molecules produced by each TF (Equation (21) in Supplementary Methods) using `transfactor`. We then model the estimated counts using a negative binomial generalized linear model as implemented in `glmGamPoi` [Ahlmann-Eltze and Huber, 2020]. The design matrix corresponds to the conditions of interest, i.e., cell type groups or pseudotime bins, and inference is carried out using *F* -tests, using the nominal *F* -distribution to compute *p*-values. The `viper` and `AUCell` methods do not assign molecules to TFs, but rather calculate enrichment or activity scores that are not counts. We therefore fit Gaussian linear models as implemented in `limma` using empirical Bayes shrinkage of the variance parameters. This approach was also adopted in Holland et al. [2020]. Here also, the design matrix corresponds to the conditions of interest, and inference is carried out using *F* -tests.

#### 2.4.3 Gamma-model-based simulation framework

In this section, we describe a Gamma-model-based simulation framework. The data-generating mechanism follows the assumptions of our proposed statistical model; `transfactor` is therefore expected to perform well in this scenario. Specifically, the simulation is based on a Gamma distribution for the TF activities 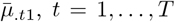, across all transcription factors in cell type 1, chosen, without loss of generality, as the reference cell type. It requires as input a shape parameter (*h* = 1 by default) and a scale parameter (*s* = 2 by default) for the Gamma distribution. Given these parameters and a gene regulatory network A, a dataset is simulated as follows.

1. Simulate TF activity for the first (reference) cell type: 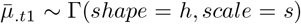, independently across all TFs *t* = 1,…, *T*.
2. For a subset of TFs chosen to be differentially active across conditions, simulate fold-changes between each cell type *c* and the reference cell type (*c* = 1) from a log-normal (*lN*) distribution: *δ*_*tc*_ ∼ *lN* (*µ* = 2.5, *σ*^2^ = .3^2^), independently across cell types *c* = 2,…, *C*.
3. In order to induce both increasing as well as decreasing TF activity between cell types, randomly set sign of log-fold-changes log *δ*_*tc*_ using independent Bernoulli(1/2) random variables.
4. For each cell type *c* = 2,…, *C*, define 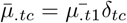.
5. Calculate 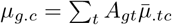, under assumptions (A1) and (A2), where 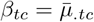 and we further assume that *α*_g*c*_ = 1.
6. Simulate gene expression measures 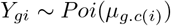.

We consider a scenario with *C* = 5 cell types and simulate gene expression measures for 50 cells of each type. The true gene regulatory network consists of *G* = 500 genes, regulated by *T* = 80 transcription factors, using only activating edges. For each gene, the number of regulating TFs are sampled from a Binomial distribution with 30 draws and success probability of 0.05. If the Binomial draw equals zero, a new draw is taken until a positive number is obtained. The set of regulating TFs for each gene are then chosen randomly. We simulate a random set of 25% of the TFs to be differentially active between cell types. In order to incorporate prior information on TF activity in `transfactor`, we let

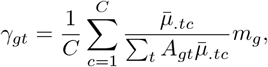

where *m*_*g*_ = 100 for all genes by default. This is then used to parameterize the prior Dirichlet distribution.

Methods are evaluated using the area under the receiver operating characteristic (ROC) curve, reflecting their capability of recovering the TFs that are differentially active between cell types.

All methods are evaluated using the true GRN, as well as GRNs where (a) edges in the true GRN have been removed; (b) edges in the true GRN have been added; (c) ‘noisy’ TFs, i.e., TFs that did not contribute to the data-generating process and regulate a random set of genes, are added.

#### 2.4.4 dyngen simulation framework

In a second simulation-based evaluation, we use the `dyngen` framework to simulate scRNA-seq data. `dyngen` is a stochastic simulation algorithm for dynamic biological processes, which simulates (nucleic acid and protein) molecular abundances at single-cell resolution and explicitly models key biological processes such as transcription, splicing, translation, and regulation. The simulation framework first constructs a gene regulatory network, including both activating as well as repressing interactions between TFs and target genes, from which molecular abundances are simulated. We can therefore use this framework to evaluate TF activity estimation, using the simulated mRNA abundances. As in the Gamma-model-based simulation framework, we benchmark several settings by adding noisy edges, removing true edges, and adding noisy TFs to the true GRN, while checking how well methods can recover the data-generating TFs. The data-generating GRN was constructed from a linear backbone in `dyngen` and consists of ∼ 600 genes regulated by 100 to 200 TFs; exact numbers vary between simulated datasets.

As prior information, we use the ‘strength’ parameter defined in `dyngen`, which is a positive number used in the data-generating scheme that approximates the strength of the regulatory interaction between a TF and its target genes. Note, however, that this prior information is only a proxy for the true strength of the edges, as several other parameters are at play in this more complex simulation scenario, and the prior information may therefore be considered more noisy, as compared to the Gamma-model-based simulation framework.

#### 2.4.5 Yeast knock-out case study

The yeast dataset was previously published in Jackson et al. [2020] and is provided in the Supplementary Material accompanying that paper. The dataset consists of 6, 829 genes and 38, 225 cells, representing 12 gene deletion mutants across 11 different environmental conditions; see Results for more details. The gene regulatory networks estimated as part of the original publication are also provided in the Supplementary Material. We use the global network aggregated across all environmental conditions (i.e., signed network.tsv file in the Supplementary Material of the original paper) to estimate TF activity. The network consists of 2, 445 genes and 129 TFs, with 1, 052 repressing and 11, 176 activating edges.

TF activity estimation was performed using the Poisson model allowing for repressions, but not enforcing sparsity. The heatmap of TF activity is created by standardizing estimated TF activities 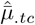 for each TF separately to have a mean of zero and a variance of one across cells. The dendrograms are created using the Ward clustering method [Ward, 1963] based on Euclidean distances, and the heatmap is visualized using the pheatmap R package. Clustering is performed by ‘cutting’ the dendrogram at a certain height in order to obtain the prespecified number of clusters.

For the diauxic shift analysis, we verify the activity of TFs identified in Murphy et al. [2015] as being ‘late-induced’, ‘early-induced’, or ‘early-repressed’ according tof analysis, we use the review paper by the TF enrichment analysis performed by the original authors. For the nitrogen metabolism analysis, we use the review paper by Conrad et al. [2014] to identify four TFs that should be upregulated during a lack of preferred nitrogen sources.

To investigate the stability of the results with respect to the cells being sequenced, we perform a bootstrap analysis, where we randomly resample cells with replacement to reach an equally large dataset, but where the number of cells for each environmental condition or genotype may vary across bootstrap samples. TF activities are then estimated on each of 30 bootstrapped dataset using the same procedure as on the full dataset.

To incorporate confidence for each edge in the GRN, we use the inferelator method that was also used in [Jackson et al.2020]. While the global network in Jackson et al. [2020] was derived using the AMuSR implementation, here we estimate the networks using the Bayesian best subset regression (BBSR) approach implemented in inferelator as it was more readily available for our purpose. We use inferelator’s built-in implementation to bootstrap cells and re-estimate the GRN five times. The accompanying confidence scores for each edge present in the original GRN are leveraged as prior information when estimating TF activity using the hierarchical Poisson model with known Dirichlet prior, where the alphaScale parameter is set to 1, i.e., *S* = 1 following the notation in the paragraph on ‘Scaling *γ*_*gt*_’ in Supplementary Methods. This means that we are giving equal weight to the information provided by the prior as well as the data.

Finally, we want to compare `transfactor` to `viper` and `AUCell` on equal grounds, as `transfactor` may be considered to have an a priori advantage by using the groupings of cells via the design matrix, which is not used for `viper` and `AUCell` since they estimate TF activity per cell separately. To avoid a possible competititve advantage, we estimate TF activity by grouping sets of at most 10 cells within each environmental condition. For `transfactor`, we use these cell sets to specify the design matrix. For `viper` and `AUCell`, we sum the counts for each cell within each set, and use these sums to estimate TF activity. Such an analysis allows an unbiased comparison between `transfactor`, which is supervised via the design matrix, and `AUCell` and `viper`, which are unsupervised in that sense.

#### 2.4.6 Olfactory epithelium case study

The olfactory epithelium dataset was previously published in Brann et al. [2020]. Here, we only focus on the cells that were assigned to the neuronal lineage. Cells belonging to this lineage were selected using the cell assignment procedure in tradeSeq, based on a trajectory fitted using `Slingshot` [Van den Berge et al., 2020, Street et al., 2018]. After subsetting, the dataset consists of 14, 618 genes and 6, 810 cells. The gene regulatory network was estimated using `grnboost2`, implemented in `SCENIC` 0.10.2 [Aibar et al., 2017]. The estimated GRN contains 7, 863 genes, 262 transcription factors, and 25, 896 edges. The median number of genes regulated by a TF is 27.

TF activity is estimated using the Poisson model with sparse initialization. We partition the cells into 20 bins, each containing ∼ 340 cells, according to the cells’ pseudotime inferred by `Slingshot`. We use this binning factor to construct a design matrix for estimating TF activity with `transfactor`.

The contribution of a TF to a cell’s overall gene expression is calculated (using the formula in Equation (21) in Supplementary Methods) and scaled by the cell’s total gene expression measure, i.e.,

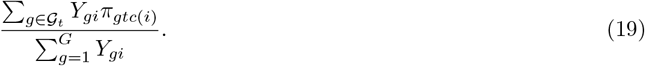

For gene set enrichment analysis on the three TF groupings as obtained in Results, we select, for each TF, the genes for which that TF is found to produce most of the molecules as compared to the other TFs regulating that gene (i.e., the genes *g* for which the TF has the highest *π*_*gtc*_ among all *t*). This set of genes for all TFs in a group is then used for gene set enrichment analysis using gProfiler [Raudvere et al., 2019], with Ensembl annotation v102, as can be accessed using https://biit.cs.ut.ee/gprofiler_archive3/e102_eg49_p15/gost.

For dimensionality reduction, we perform principal component analysis (PCA) using scater [McCarthy et al., 2017] to derive the top 10 principal components on the output of each method. Since our method outputs counts, we log-transform them before PCA. The top 10 PCs are then used as input for uniform manifold approximation and projection (UMAP) dimensionality reduction [McInnes et al., 2018] with parameter min dist=0.8.

## 3. Results

### 3.1 Simulation studies

#### 3.1.1 Gamma-model-based simulation

##### Estimation of TF activity parameters

In the gamma-model-based simulation framework, we know the true TF activities and their corresponding contributions to gene expression. Hence, we can evaluate how close the estimates from `transfactor` are to the true parameter values using the mean squared error (MSE) (Supplementary Feigure 1). Note that this is not possible for `viper` and `AUCell`, as their output is on a different scale and therefore non-interpretable in terms of TF activity in the way we simulate it. The `poisson` and dirMultEBayes models perform similarly in terms of MSE across all conditions, while providing prior information using the dirMult model results in more accurate estimation, especially when the GRN contains many false edges and truly non-active TFs. For all models, the sparse initialization has no influence on the MSE, likely because the truly non-active TFs have very low 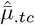 either way; therefore, setting them to zero has negligible impact on the MSE, which is dominated by truly active TFs with high 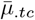.

##### Detection of differential TF activity

Next, we turn to assessing the performance of TF activity inference methods in terms of their ability to detect differentially active TFs. We use the area under the receiver operating characteristic curve (AUC) as a performance measure; a high AUC corresponds to a high sensitivity and specificity for identifying the TFs truly involved in the system under study. We first discuss the scenario where we know and use the true data-generating GRN (middle panel in top row of Figure 1). In this scenario, using a sparse initialization of the EM algorithm provides no benefits, as expected. While all methods perform generally well, the `poisson` model performs best, as does `AUCell`. Note that, while it may seem counterintuitive at first that the dirMult model performs somewhat worse, this can be explained by the fact that the prior distribution is shared across cell types, effectively leading to moderation of the TF activity estimates by the prior distribution across cell types. Since the performance measure used here relates to differential TF activity, this moderation can lead to an attenuation of differences in TF activity, hence loss of power, whenever the true TF activities are different across cell types.

**Figure 1:**
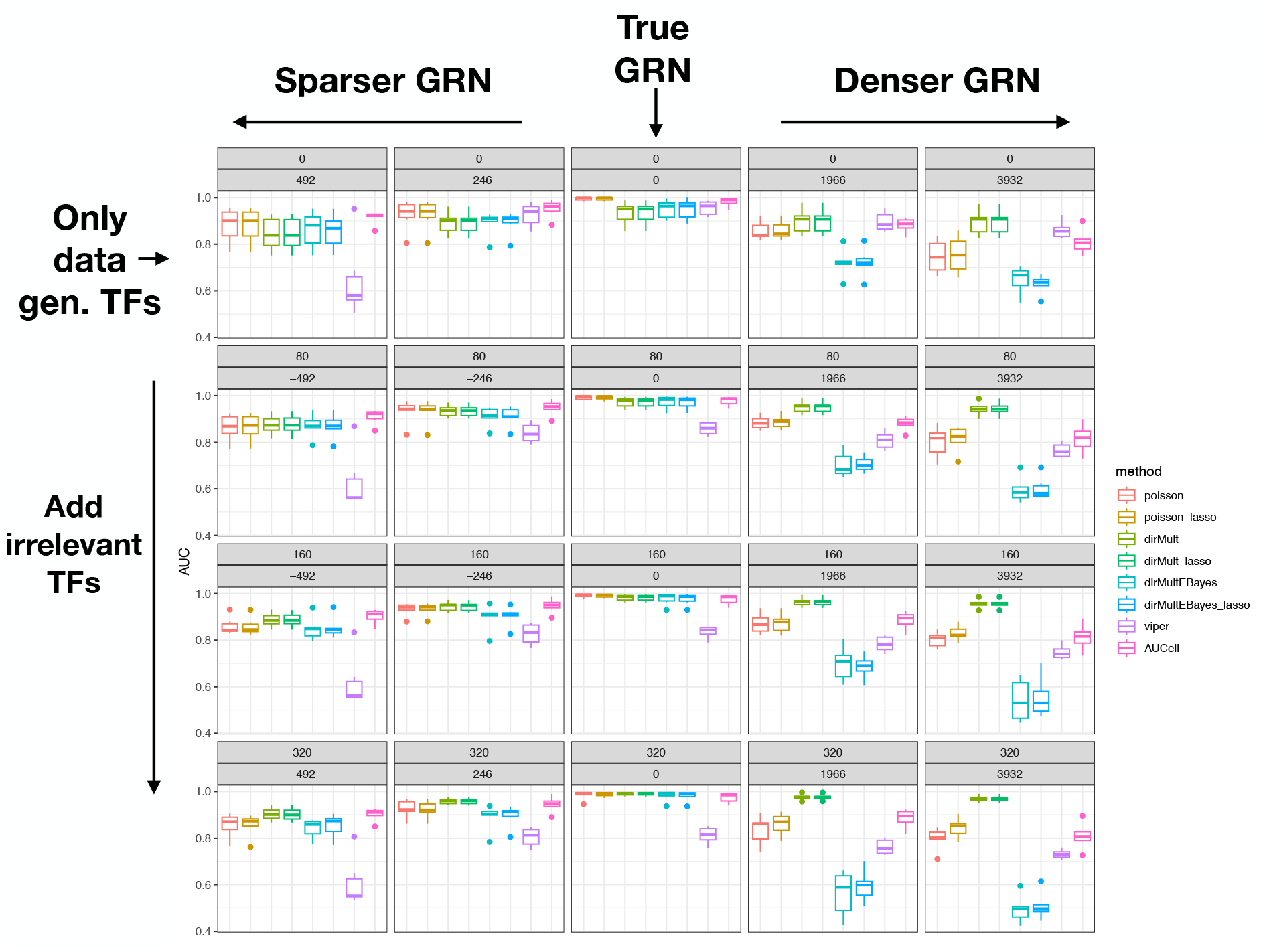
Gamma-model-based simulation study results. Eight methods were evaluated on each of six simulated datasets. For each method and dataset, the area under the receiver operating characteristic curve for detecting the TFs that are differentially active between cell types was computed. For each method, the six AUCs were summarized using a boxplot. The different panels represent different GRNs used as input to each method. The first row corresponds to a GRN where only TFs that were involved in the data-generating process are used. For rows 2–4, respectively, 80, 160, and 320 ‘irrelevant’ TFs (i.e., TFs that do not contribute to the data-generating process and regulate a random set of genes) were added to the GRN. The different columns represent removing or adding edges to the true GRN. In the middle column, we use the true GRN. Moving to the left (right) of the middle column, we remove (add) edges to the true GRN. Panel subtitles indicate the number of irrelevant TFs added to the GRN and edges added/removed from the GRN.

Next, we evaluate a more realistic scenario, where we do not know the true GRN (see Methods for details). First, we consider the scenario where edges are either removed from or added to the true GRN (i.e., the first row in Figure 1). As edges are removed, all methods drop in performance, but both the `poisson` model and `AUCell` still retain good performance. `viper` is the only method for which performance declines severely. As more false positive edges are added to the GRN, there is a markedly larger drop in performance for most methods. As could be expected, the dirMult methods are capable of using the prior information to prioritize true edges in the GRN and thus perform best, while the `dirMultEBayes` methods perform worst. Among the competing methods, `viper` seems more robust to false positive edges than `AUCell`.

We also consider the scenario where ‘irrelevant’ TFs are added to the GRN. In general, we see that sparse initialization can become beneficial here as, indeed, the irrelevant TFs are identified and their activities are shrunken to zero. This is especially apparent in the setting where the GRN has both many irrelevant TFs and many false positive edges, i.e., the bottom right panels of Figure 1. Note that this is likely also the setting that corresponds most closely to a real data analysis, when using a GRN that could, for example, be downloaded from a database and not necessarily be appropriate for the specific context. The dirMult methods perform consistently very well, since they are able to use prior information to prioritize true edges and data-generating TFs. In settings where the GRN can be expected to contain a lot of noise, but there is no prior information available, the `poisson` method with sparse initialization performs well in general.

For the differential activity (DA) analysis based on estimated TF activities from `transfactor`, we have made use of count models as implemented in `glmGamPoi` [Ahlmann-Eltze and Huber, 2020], while we have used `limma` [Smyth, 2004] for `viper` and `AUCell`, as their output cannot be considered to be count data. In order to prevent such differences in post-processing from confounding the comparison of `transfactor`, `AUCell`, and `viper`, we also provide a version of Figure where `limma` is used to perform differential activity analysis for all methods (Supplementary Feigure 2) and the voom transformation approach is applied for all `transfactor` methods [Law et al., 2014]. While in general the performance of `transfactor` is lower, most of the results are qualitatively similar. The dirMult model, however, seems to specifically suffer from the change in DA analysis methodology. Finally, `AUCell` performs better than `transfactor` when the GRN is missing true edges.

Taken together, these results suggest that, if reliable prior information is available, it is beneficial, as expected, to use the dirMult model. However, in real data settings, we generally don’t have as accurate prior information as we have here. When the GRN can be expected to contain both false positive edges and TFs that are not involved in the process being studied, the sparse initialization step helps, and the `poisson` model could be a good choice. In cases where we expect the GRN to be missing true edges, `AUCell` also performs well.

3.1.2 `dyngen` simulation

We also evaluate TF activity estimation using the `dyngen` simulation framework [Cannoodt et al., 2020]. We simulate six datasets, each of which can be represented by a trajectory consisting of a single lineage in a low-dimensional embedding (see Supplementary Feigure 3a, for an example). Note that, since `dyngen` uses a gene regulatory network to simulate the data (Supplementary Feigure 3c–d), we know the true gene regulatory network that is driving the development represented by this trajectory. Each dataset consists of ∼ 700 cells, for each of which we are observing 100–200 TFs, ∼ 500 target genes, and ∼ 100 housekeeping genes (that are not regulated by any TFs). The GRN used to generate the data includes activating as well as repressing edges, additionally allowing us to benchmark our approach to account for repressions in the GRN. The prior information in this setting is more noisy as compared to the Gamma-model-based simulation framework, due to a more complex simulation scenario.

First, evaluating all methods using the true GRN as input (middle panel in top row of Figure 2), it is clear that `transfactor` outperforms competing methods `viper` and `AUCell`. As for the gamma-model-based simulation of Section 3.1.1, sparse initialization has no apparent impact on the performance of the `transfactor` methods if the TF activities are in fact not sparse. Accounting for repressions in addition to the sparse initialization does not seem to increase performance in this setting. When true edges are removed from the GRN (left panels in top row of Figure 2), all methods similarly drop in performance, while `AUCell` remains rather stable. As false edges are added to the GRN (right panels in top row of Figure 2), using the available prior information on the strength of TF-gene interactions from the `dyngen` simulation framework provides stability to the results, and accounting for repressions can become more beneficial. Overall, our `transfactor` methods perform better than `AUCell` and `viper`.

**Figure 2:**
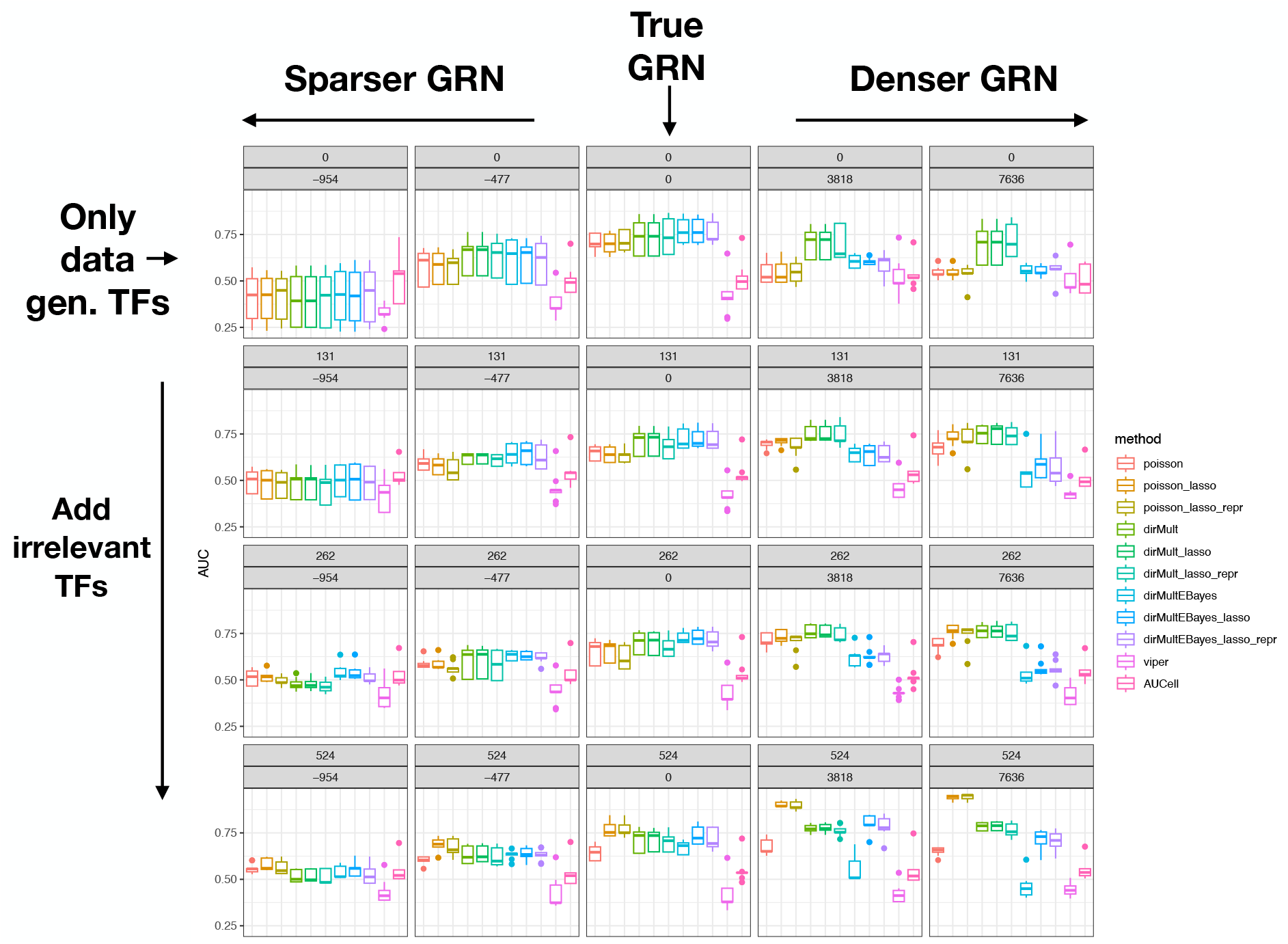
dyngen simulation study results. Eleven methods were evaluated on each of six simulated datasets. For each method and dataset, the area under the receiver operating characteristic curve for detecting the TFs that are differentially active between the cell types was computed. For each method, the six AUCs were summarized using a boxplot. The different panels represent different GRNs used as input to each method. The first row corresponds to a GRN where only TFs that were involved in the data-generating process are used. For rows 2–4, respectively, 131, 262, and 524 ‘irrelevant’ TFs (i.e., TFs that do not contribute to the data-generating process and regulate a random set of genes) were added to the GRN. The different columns represent removing or adding edges to the true GRN. In the middle column, we use the true GRN. Moving to the left (right) of the middle column, we remove (add) edges to the true GRN. Panel subtitles indicate the number of irrelevant TFs added to the GRN and edges added/removed from the GRN.

As more irrelevant TFs are added to the GRN (rows 2–4 in Figure 2), sparse initialization results in larger gains, especially when many false positive edges are present in the GRN. Note that the sparse initialization has no real impact on the dirMult model, since it is overridden by the prior information. As edges are removed from the true GRN as well as a large number of irrelevant TFs are added, the `poisson`, dirMult, and `dirMultEBayes` methods start performing similarly.

Note that when many false positive edges are added to the GRN as well as many irrelevant TFs, the dirMult method is inferior to the `poisson` method with sparse initialization. This is due to multiple reasons. First, the prior includes information on edges from irrelevant TFs in the GRN, which overrides the sparse initialization (see Methods), hence losing its performance benefit. It is thus more appropriate to compare the dirMult methods to the `poisson` method without sparse initialization. Second, the prior parameters for noisy and true edges are often of similar magnitude, due to the strength parameters provided by `dyngen`. The effect observed is therefore partially the result of a less discriminative prior; see Supplementary Feigure 4 for the results when using a prior that is always higher for true vs. false edges. Third, the prior parameters here are only a proxy for their true value, while in the Gamma-model-based simulation study their true value was known.

We again investigate performance for all methods when exclusively using `limma`(-voom) for differential activity analysis. In this simulation study, we notice little influence of the downstream DA approach on methods’ performance (Supplementary Feigure 5).

In summary, these results largely confirm those obtained with the Gamma-model-based simulation framework, but additionally show that accounting for true repressing interactions can be beneficial. If reliable prior information on the interaction strength is available, it would be a good idea to use it, however, noisy prior information can also be detrimental, especially if it overrides the sparse initialization step that can be quite powerful. If such information is not available, methods based on the Poisson and empirical Bayes hierarchical Poisson models are powerful alternatives.

### 3.2 Case studies

#### 3.2.1 Yeast knock-out dataset

We analyze a scRNA-seq dataset on budding yeast (*Saccharomyces cerevisiae*) [Jackson et al., 2020]. The dataset encompasses 12 transcriptionally-barcoded gene deletion mutants, each investigated in 11 different environmental conditions, comprising a total 38, 225 single cells. As noted by the authors, the different environmental conditions represent the largest source of variability in the dataset, with the effects of mutations hardly noticeable in reduced-dimensional space [Jackson et al., 2020]. The environmental conditions are summarized in Table 1. The original publication used the diversity of transcriptional states, in combination with known regulations, to construct a global gene regulatory network across mutants and environmental conditions, consisting of 2, 445 genes and 129 TFs, including 1, 052 repressing and 11, 176 activating edges. We use this GRN to estimate TF activity, accounting for repressions, using our Poisson model. The design matrix accommodates a possible interaction of genotype (gene deletions) and environmental condition. Given the careful construction of the GRN using a range of transcriptional states and prior information relevant to the dataset, we do not enforce sparsity.

**Table 1:** Yeast knock-out case study. Environmental conditions.

| Condition | Nutrient characteristics |
| --- | --- |
| YPD | Yeast Extract, Peptone, D-Glucose. A normal growth condition, where glucose is the main energy source through glycolysis, releasing ethanol. |
| YPDDiauxic | YPD (Harvested after Post-Diauxic Shift). When glucose becomes limiting, the cells enter diauxic shift characterized by decreased growth rate and by switching metabolism from glycolysis to aerobic utilization of ethanol. |
| YPDRapa | YPD + 200 ng/mL Rapamycin. Rapamycin is a potent inhibitor of TOR and inhibits proliferation of yeast cells. |
| YPEtOH | Yeast Extract, Peptone, Ethanol. Like YPD, except that D-Glucose is replaced by ethanol, therefore switching the main carbon source to ethanol. |
| MIN_Gluc | Minimal media with Glucose as main carbon source. Ammonium sulfate is main nitrogen source. |
| MIN_EtOH | Minimal media with ethanol as main carbon source. Ammonium sulfate is main nitrogen source. |
| NLIM_GLN | Nitrogen limited medium with Glutamin. Nitrogen source is therefore Glutamin. |
| NLIM_NH4 | Nitrogen limited medium with Ammonium Sulfate. Nitrogen source is therefore Ammonium Sulfate. |
| NLIM_PRO | Nitrogen limited medium with Proline. Nitrogen source is therefore Proline. |
| NLIM_UREA | Nitrogen limited medium with urea. Nitrogen source is therefore urea. |
| CStarve | Carbon-deprived medium. No Carbon source available. |

The heatmap in Figure 3a displays the estimated mean TF activities for each mutant in each environmental condition for each of the 129 TFs from the GRN, where, for each TF, activities are standardized to have zero mean and unit variance across cells. As in the original publication, samples cluster mainly by environmental condition. The estimated TF activities are useful summaries of the data, as indicated by their effectiveness at recovering the environmental conditions, here, by hierarchical clustering followed by cutting the dendrogram at a certain height such that the number of clusters corresponds to the number of environmental conditions. The adjusted Rand index (ARI) of such a TF-activity-based clustering corresponds to 0.83, as compared to an ARI of only 0.53 when clustering samples according to the average expression of those same TFs (here, the ARI measures the similarity between the clusterings of samples based on TF activity or expression measures and the reference grouping of samples based on environmental condition). The heatmap of the standardized estimated TF activities highlights some remarkable patterns, in particular, the condition-specific higher activity of TFs and transcriptional states for some conditions that seem to be governed by a limited number of most active TFs, while other TFs show relatively moderate to low activities, across most mutants. Examples are the YPD and CStarve conditions, amongt others. In the YPD condition, where all nutrients are available, a group of seven TFs (*IFH1, SFP1, FKH2, HMS1, HCM1, URC2*, and *MIG1*) cluster together as being most active in that condition (purple box in Figure 3a). This group of TFs includes *IFH1*, a TF known to regulate ribosomal protein (RP) genes and recruited in optimal growth conditions [Schawalder et al., 2004]. *SFP1* is known to be essential for normal yeast cell growth [Blumberg and 1991], and also regulates RP genes [Jorgensen et al., 2004], while also contributing to G2/M transitions in the mitotic cell cycle [Xu and Norris, 1998]. *FKH2* is similarly involved in cell cycle dynamics, being a known activator of replication origins, and regulates expression of G2/M phase genes. Not much is known about *HMS1*, however, overexpression of the TF results in hyperfilamentous growth, hence it could be required for yeast cell growth [Lorenz and Heitman, 1998]. *HCM1* drives S-phase activation of genes involved in chromosome segregation [Pramila et al., 2006]. is involved in glucose repression, whereby cells grown on glucose repress the expression of a large number of genes that are required for the metabolism of alternative carbon sources [Carlson, 1999]. Finally, there does not seem to be much known about *URC2*, a TF that is possibly involved in uracil catabolism [Andersen et al., 2008], a function that would align with those of other TFs in this group.

**Figure 3:**
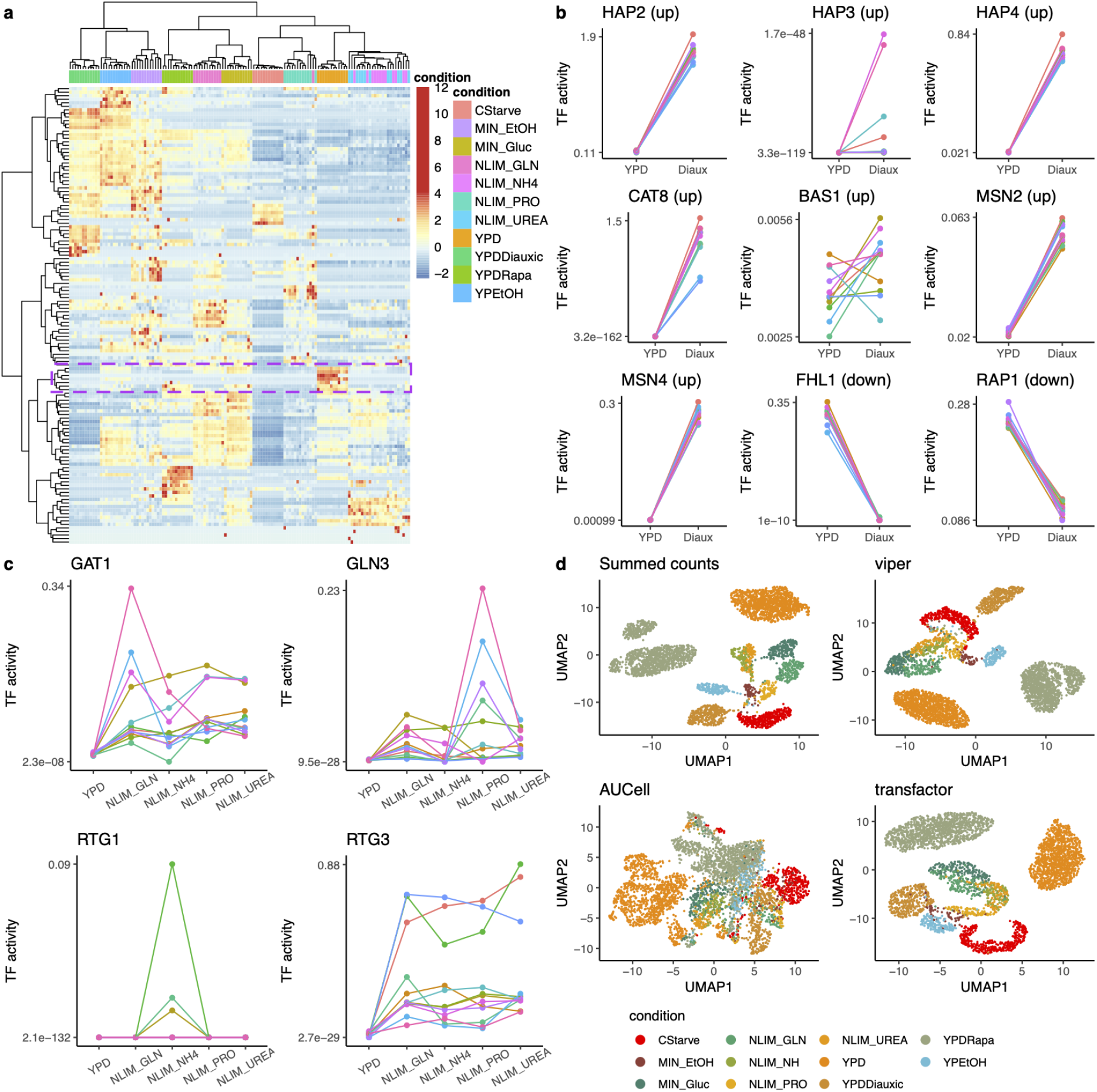
Yeast knock-out case study. (a) Heatmap of estimated TF activities 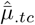 from `transfactor`, where rows correspond to TFs and columns to cell groups based on condition and genotype. The pseudocolor correspond to TF activity, standardized to have zero mean and unit variance across cell groups. (b) TF activity estimates from `transfactor` for nine TFs involved in the diauxic shift. Each color represents a different gene deletion mutant. (c) TF activity estimates from `transfactor` for four TFs involved in nitrogen metabolism. Each color represents a different gene deletion mutant. (d) UMAP dimensionality reduction plots for the gene expression counts from aggregated cells (‘Summed counts’, see Methods) and corresponding TF activity estimates using `viper`, `AUCell`, and `transfactor`. Cells are colored according to environmental condition.

Aside from the initial exploratory data analysis, we also verify if `transfactor` can capture regulators known in the literature to be associated with changes caused by particular environmental conditions. Murphy et al. [2015] study protein abundances during the diauxic shift using a proteomics time series experiment. They identify protein lists that are either induced or repressed, early or late in the diauxic shift, and subsequently use the YEASTRACT transcription factor enrichment tool [Teixeira et al., 2023] to assess which transcription factors might drive the observed differences in protein abundance. This procedure identifies a total of nine TFs, seven of which should be upregulated in the diauxic shift and two which should be downregulated. Our estimated TF activities confirm the findings of the proteomic study, and show very good agreement of up- and downregulation of TFs in the diauxic shift, as compared to the control (YPD) condition (Figure 3b). One notable exception is the *BAS1* TF, where the up- or downregulation is variable across different mutants. For *HAP3*, we find very low TF activity in both conditions.

The dataset also includes four environmental conditions that are related to nitrogen limitation, where the preferred nitrogen source (ammonium sulfate) is replaced by alternative sources. Conrad et al. [2014] review nutrient sensing and signaling in yeast. They discuss that, in the absence of a preferred nitrogen source, four TFs (*RTG1, RTG3, GLN3*, and *GAT1*) are activated, while in the presence of a preferred nitrogen source these are deactivated through (hyper)phosphorylation. We therefore compare TF activity of all four nitrogen-limitation conditions with the YPD condition, and indeed detect increased TF activity in all nitrogen conditions for most of these TFs, confirming what is known in the literature (Figure 3c).

While confirming known biology is important, the results should also be stable with respect to the cells being sequenced. We therefore performed a bootstrap analysis, by randomly resampling cells with replacement to get an equally large dataset, and re-estimating TF activity to see if our conclusions still stand. The changes in TF activity for both the diauxic shift as well as the nitrogen limitation TFs are stable across bootstrap samples (Supplementary Feigures 6–7).

Furthermore, we assess the stability of the results when incorporating information on the confidence of the edges in the GRN. To this end, we re-estimate the GRN across five different bootstrap samples using the inferelator method (see Methods), which the authors of the study used to infer their GRN, and leverage the confidence scores (i.e., the probability each edge is present across the bootstrap samples) for each edge as prior information in the TF activity estimation using the hierarchical Poisson model. Incorporating the uncertainty on GRN edges results in TF activities that equally well cluster the samples according to environmental condition, with an ARI of 0.82. The biological findings on the diauxic shift and nitrogen metabolism are also confirmed, with the exception that the *BAS1* TF is now less active in the diauxic condition as compared to the control (Supplementary Feigure 8).

We also compare `transfactor` with `viper` and `AUCell` in terms of recovering the biological structure (i.e., the different environmental conditions) of the data upon TF activity estimation. Both `viper` and `AUCell` are unsupervised methods, assessing TF activity for each cell separately, while `transfactor` estimates TF activity separately within each of a predefined set of cell types. To be able to compare all methods on an equal footing, we first group cells in random sets of 10 within each environmental condition, and use this grouping as design matrix for `transfactor`, while for `viper` and `AUCell` we add counts for these sets of cells and then assess TF activity estimation. This analysis strategy ensures that all methods use the same information and avoids a priori competitive advantages. Figure 3d shows UMAP representations of the counts and corresponding TF activity estimates from all methods. While `viper` and `transfactor` recover much of the biological variability, `AUCell` fails at clearly separating many environmental conditions. This is also reflected in the ARI, which is 0.32 for both `viper` and `transfactor`, and 0.21 for `AUCell`.

Finally, one may wonder whether the results obtained in this case study may have similarly been obtained without estimating TF activity, and solely considering TF expression instead. We have already seen that TF activity estimates better reflect the biological structure of the dataset, as noted by the increased ARI when comparing activity-based (vs. expression-based) clusters to environmental conditions. The confirmation of increased and decreased TF activity in the diauxic shift condition are much less clear when looking at TF expression rather than TF activity. Indeed, the expression of *BAS1* and *MSN2* conflicts with what is known in the literature. Further, the two TFs that should decrease in activity (*FHL1* and *RAP1*) now no longer consistently do so in terms of expression (Supplementary Feigure 9a). Similarly, results on the nitrogen limitation condition show less agreement with the literature when looking at TF expression as compared to TF activity (Supplementary Feigure 9b).

In summary, this case study shows that TF activity estimates are relevant and useful, allowing novel insight in terms of TF regulation. It additionally demonstrates the flexibility of our method to handle different situations with varying levels of information on the GRN.

#### 3.2.2 Olfactory epithelium dataset

The olfactory epithelium (OE) is responsible for the perception of smell and sending these molecular signals to the brain. It also serves as a barrier between the outer world and the brain, requiring neuronal cells to be continuously replaced through a source population of progenitor cells. In Brann et al. [2020], the OE is chemically injured to trigger regeneration and repair of the tissue through activation of stem cells known as horizontal basal cells (HBC). To develop into neuronal cells, HBCs first differentiate into globose basal cells (GBC), which mitotically divide toensure continuous neuronal regeneration [Fletcher et al.,, 2017]. These GBCs in turn develop into immature neurons and finally mature neurons [Fletcher et al., 2017, Gadye et al., 2017]. The trajectory is shown in Figure 4a.

**Figure 4:**
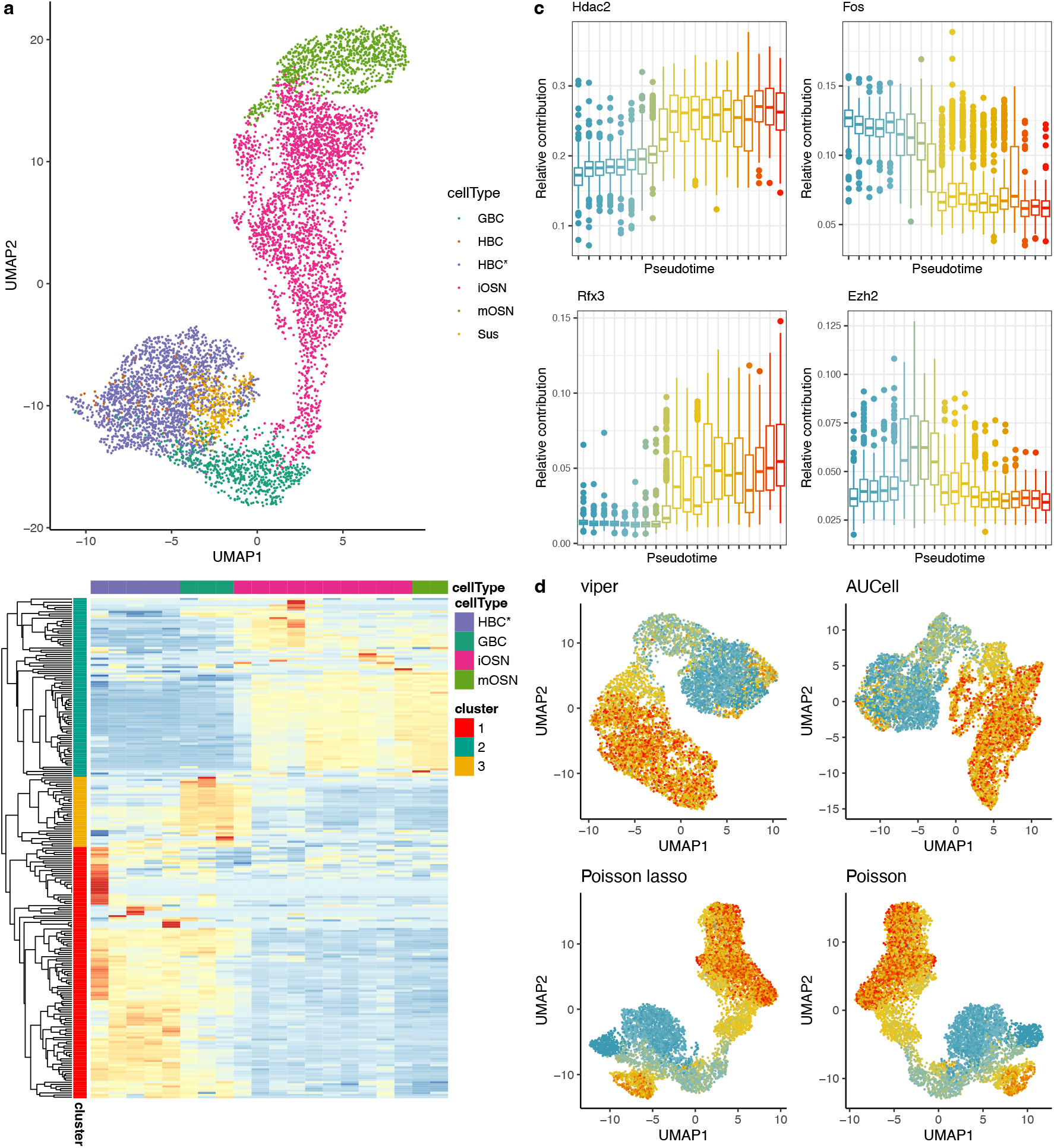
Olfactory epithelium case study. (a) UMAP dimensionality reduction of the gene expression data, colored according to cell type. A clear differentiation path from activated horizontal basal cells (HBC*) to neuronal cells is observed. (b) Heatmap of estimated TF activities 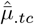from `transfactor` for the 235 TFs found to be involved in the neuronal lineage. Each row represents a TF and the different columns represent 20 cell groups of approximately equal size, binned according to their pseudotime values. The pseudocolor correspond to TF activity, standardized to have zero mean and unit variance across bins. Three clusters are identified based on their activity patterns. (c) Activity patterns of the top 4 transcription factors, where the top list was derived using the scaled Euclidean distance measure (see Equation (4) in Methods). The y-axis represents the contribution of a TF to the entire observed gene expression in each cell (Equation (19)), e.g., if the contribution equals 0.1, then we estimate that that TF was involved in the production of 10% of all molecules present in the cell. The x-axis represents the 20 pseudotime bins. Note that panels are on different y-axis scales. (d) UMAP reduced-dimensionality plots based on estimated TF activity for four methods. Cells are colored according to their pseudotime bin, from dark blue (first bin) to red (last bin). Note that panels are on different scales. co-regulated modules. Alternatively, community detection algorithms (e.g., [Li et al., 2020]) could be used.

TF activity is estimated using the `transfactor` `poisson` model with sparse initialization. Out of a total of 262 TFs that are included in the GRN estimated by `SCENIC` [Aibar et al., 2017], our model finds that 27 do not actively contribute to the process, i.e., zero molecules are associated with these TFs. A heatmap of the standardized activities of the contributing TFs shows three distinct groups of TFs which can be discriminated based on their activity patterns (Figure 4b). These groups are respectively most active in the HBC, GBC, and neuronal stages, recapitulating the biological knowledge on these three previously-identified cell types [Fletcher et al., 2017, Gadye et al., 2017]. The relevance of the grouping is confirmed by gene set enrichment analysis on the genes found to be regulated by these TF groups, which identifies significant biological processes involved in tissue development, cell cycle, and cell projection for, respectively, the HBC, GBC, and neuronal groups. The top enriched gene sets for each group are provided in Supplementary Tables 1–3.

High-throughput studies typically focus on differential expression analysis to investigate gene regulation. Here, we identify differentially expressed genes along the neuronal lineage using tradeSeq [Van den Berge et al., 2020] and we find 6, 977 DE genes at a 5% nominal false discovery rate (FDR) level; a long list of genes that is cumbersome to interpret. Often, one tries to shorten the list of DE genes by imposing a fold-change cut-off (e.g., McCarthy and Smyth[2009b]). However, testing against a fold-change cut-off of 2, still results in 4, 752 DE genes. In comparison, again using tradeSeq, we find that 209 TFs are differentially active along the neuronal lineage, 135 of which are still differentially active after testing against the same fold-change threshold, leading to a much more parsimonious interpretation of the dataset.

We use a scaled Euclidean distance-based ranking of transcription factors (see Methods) to order them according to their contribution to the observed differences in gene expression. Here, we focus on the top 4 transcription factors, which collectively explain 28% of the total scaled Euclidean distance across all TFs. The TF activities, scaled by the the total activity across all TFs, are shown in Figure 4c. The top transcription factor, *Hdac2*, is a member of the Hdac family, which is interestingly known to be essential in the response of neurons following injury [Cho and Cavalli, 2014]. *Rfx3* is required for cilia formation in neurogenesis [Choksi et al., 2014, is known to be involved in the injury, activation, and differentiation of stem cells [Okada et al., 1996,, 2015]. *Fos* and Weems, 1999], and is indeed most active at the early stages of development in the lineage. *Ezh2* is most active at the GBC stage; previous work confirms that *Ezh2* protein activity is indeed found to be highest in the GBC stage [Goldstein et al., 2018]. In summary, the literature confirms that our top transcription factors are involved in olfactory neuron maturation and regeneration upon injury.

On a larger scale, we check how well our method, but also `viper` and `AUCell`, are able to recover the global structure of the data following TF activity estimation. We therefore run `viper` and `AUCell` on all cells using the same GRN estimated by `SCENIC`. Note, however, that in this analysis our method makes explicit use of the grouping of cells into 20 equally-sized bins according to pseudotime, while `viper` and `AUCell` are completely unsupervised, and we therefore expect our method to already have a competitive advantage to recover the structure defined by the cell groupings. We perform dimensionality reduction on the output of each method, by using the top 10 principal components as input to UMAP [McInnes et al., 2018]. The global structure is generally recovered in reduced-dimensional space for all methods, where a differentiation path of cells corresponding to the pseudotime of the trajectory can be observed (Figure 4d).

Finally, we investigate the robustness of our results to an analysis where we do not use the sparse initialization step. All four top TFs discussed previously are still the top four. In general, the rankings, especially the top rankings, are highly correlated. Performing the same dimensionality reduction on the non-sparse output again leads to recovering the global structure of the data (Figure 4d).

## 4 Discussion

We have proposed the `transfactor` method to infer transcription factor activity by deconvolving transcription factor-specific gene expression measures from observed overall gene expression measures, based on a Poisson model for read counts. A hierarchical Bayes version of the model allows the incorporation of prior information on TF activity. The method is implemented in the open-source software package `transfactor` available at https://github.com/koenvandenberge/transfactor.

The `transfactor` method relies on a gene regulatory network as input, and we have found the GRN to have a large impact on downstream results. However, it is currently unclear how to assess whether a GRN, either obtained from databases or estimated empirically from the data, is appropriate for a specific dataset. For example, one would perhaps be able to check this by examining whether co-regulated genes are indeed correlated in terms of their measured gene expression.

Using TF activity instead of gene expression in downstream analyses has proven to be useful in multiple ways. By substantially reducing the dimensionality of the dataset, use of TF activities should increase the signal-to-noise ratio and allow for easier biological interpretation. In our case studies, TF activities have shown to be better than gene expression measures at grouping cells with respect to their biological condition, and the obtained changes in transcription factor activity match what is known from the literature.

The estimated TF activities allow one to prioritize TFs based on their estimated contributions to observed gene expression changes. However, a comparison of TF activity across groups of cells or along lineages of a trajectory, as done in this manuscript, effectively ignores the inherent uncertainty on the estimates. Further work should be directed to estimating the uncertainty on the estimated TF activities. For example, in transcript-level quantification of RNA-seq, where an EM algorithm is adopted to derive transcript abundances, the uncertainty is quantified by bootstrapping the data and re-running the EM algorithm [Bray et al., 2016, Patro et al., 2017]. Other approaches such as described in Louis [1982] allow for uncertainty estimation to be embedded in the framework of the EM algorithm [Bray et al., 2016, Patro et al., 2017], however, significantly increasing its computational burden.

One of the core assumptions of our model is that, if a TF is active, it is so across all of its target genes. This is surely a simplification of reality, where transcription factors may vary in their effective DNA binding according to specific sequence preferences and therefore between different target genes. An interesting direction would be to weigh the edges of the GRN using models that are capable of computationally estimating binding probabilities to specific target sequences (in this case, gene promotors), such as the DeepBind or BPNet models [Alipanahi et al., 2015, Avsec et al., 2021]. In addition, integration with other data modalities, such as the single-cell assay for transposase-accessible chromatin (scATAC-seq), could be relevant in order to detect plausible binding regions and use TF-specific binding probabilities for each of these regions. Finally, one may also consider a different parameterization of the statistical model, e.g., no longer the multiplicative model that we propose here, but a more flexible parameterization.

The sparse initialization step of the EM algorithm relies on an equal partitioning of molecules to the TFs that could have produced them. This leaves the sparse initialization step vulnerable to errors, possibly leading to both false positives and false negatives. To avoid this, one could first run the EM algorithm for a few iterations (using the same starting values as usual), after which the obtained estimates could be used in the sparse initialization step. The subsequent sparse solution can then be used as input to the following iterations.

While in our manuscript we have focused on exploring the estimated TF activities as the main tool for interpreting the datasets, our method can also be used for more detailed analysis of co-regulation. For example, for each cell type, one could construct a weighted network with the same structure as the GRN A, but where the edges have weights equal to 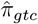. If one considers A as a bipartite graph, recent biclustering techniques (e.g., [Moran et al., 2021]) could be applied to simultaneously group genes and transcription factors together, in order to identify Acknowledgements The authors thank John Ngai for sharing the dataset that was used in the olfactory epithelium case study and for providing feedback on the obtained biological results.

