## Supplementary Material for "transfactor: Transcription factor activity estimation via probabilistic gene expression deconvolution"

### 5 Supplementary Methods

#### 5.1 EM algorithm for fitting the hierarchical Poisson model

Below, we provide details on the EM algorithm used to fit the hierarchical Poisson model. Note that, while the prior parameters $\gamma_{gt}$ may be either assumed to be known or estimated, here we describe the optimization in the context where they would have to be estimated as this is the most general case. If the prior parameters are assumed to be known, M-step II is skipped.

**Q-function.** Ignoring the term corresponding to the observed data and not including the parameter of interest $\theta = \{\pi_{gtc}, \gamma_{gt}\}$, we have

$$Q(\theta^{new}|\theta^{old}) = E\left[\log Pr(\mathbf{Z}|\mathbf{Y}, \theta^{new})|\mathbf{Y}, \theta^{old}\right]$$
$$= E\left[\sum_{i=1}^{n}\sum_{g=1}^{G}\left\{\log Y_{gi}! - \sum_{t\in\mathcal{T}_g}\left(\log Z_{gti}! - Z_{gti}\log \pi_{gtc}^{new}\right)\right\}\middle|\mathbf{Y}, \theta^{old}\right].$$

Dropping terms that are not a function of the parameter of interest,

$$Q(\theta^{new}|\theta^{old}) = \sum_{i=1}^{n}\sum_{g=1}^{G}\sum_{t\in\mathcal{T}_g} E[Z_{gti}|\mathbf{Y}, \theta^{old}]\log \pi_{gtc}^{new},$$

up to an additive constant.

**Prior.** The prior log-likelihood is

$$G(\theta) = Pr(\pi|\gamma) = \sum_{g=1}^{G}\left[\log\Gamma(\sum_{t\in\tau_g}\gamma_{gt}) + \sum_{t\in\tau_g}\{(\gamma_{gt}-1)\log \pi_{gtc} - \log\Gamma(\gamma_{gt})\}\right].$$

**E-step.** In the E-step, we calculate $Q(\theta^{new}|\theta^{old})$. This boils down to computing $E[Z_{gti}|\mathbf{Y}, \theta^{old}] = Y_{gi}\pi_{gtc}^{old} = Y_{gi}\frac{\bar{\mu}_{.tc}^{old}}{\sum_{t\in\mathcal{T}_g}\bar{\mu}_{.tc}^{old}}$, making use of Equation (9).

**M-step.** Given $Q$, estimate $\pi_{gtc}^{new}$ and $\gamma_{gt}^{new}$. This is achieved by maximizing $Q(\theta^{new}|\theta^{old}) + G(\theta^{new})$, where $G$ is the log-prior density, in order to obtain the posterior mode for $\pi_{gtc}^{new}$ and $\gamma_{gt}^{new}$ [Dempster et al., 1977].

**M-step I: Estimate $\pi_{gtc}^{new}$.** First, we estimate $\pi_{gtc}^{new}$ given $\gamma_{gt}^{old}$. By incorporating the Dirichlet prior distribution, we now have

$$Q(\theta^{new}|\theta^{old}) + G(\theta^{new}) = \sum_{i=1}^{n}\sum_{g=1}^{G}\left[\log\Gamma(\sum_{t\in\tau_g}\gamma_{gt}^{old}) + \sum_{t\in\mathcal{T}_g}\left\{\pi_{gtc}^{old}Y_{gi}\log \pi_{gtc}^{new} - \log\Gamma(\gamma_{gt}^{old}) + (\gamma_{gt}^{old}-1)\log \pi_{gtc}^{new}\right\}\right].$$

Dropping terms not involving $\pi_{gtc}^{new}$, we get

$$Q(\theta^{new}|\theta^{old}) + G(\theta^{new}) = \sum_{i=1}^{n}\sum_{g=1}^{G}\sum_{t\in\mathcal{T}_g}(\pi_{gtc}^{old}Y_{gi} + \gamma_{gt}^{old} - 1)\log \pi_{gtc}^{new}.$$

Note that we cannot maximize $Q$ directly, as we need to take into account the constraint that $\sum_{t\in\mathcal{T}_g}\pi_{gtc} = 1$. We can use Lagrange multipliers to instead maximize

$$\sum_{i=1}^{n}\sum_{g=1}^{G}\sum_{t\in\mathcal{T}_g}(\pi_{gtc}^{old}Y_{gi} + \gamma_{gt}^{old} - 1)\log \pi_{gtc}^{new} + \lambda(1 - \sum_{t\in\mathcal{T}_g}\pi_{gtc}^{new}).$$

Taking the partial derivatives with respect to $\pi$ for a particular cell type $c$, gene $g$, and transcription factor $t \in \mathcal{T}_g$,

$$\frac{\partial}{\partial \pi_{gtc}^{new}} \left( Q(\theta^{new}|\theta^{old}) + G(\theta^{new}) \right) = \sum_{i \in \mathcal{C}_c} \frac{\pi_{gtc}^{old} Y_{gi} + \gamma_{gt}^{old} - 1}{\pi_{gtc}^{new}} - \lambda,$$

where $\mathcal{C}_c$ denotes the set of cells belonging to cell type $c$. We also take the partial derivative with respect to the Lagrange multiplier $\lambda$,

$$\frac{\partial}{\partial \lambda} \left( Q(\theta^{new}|\theta^{old}) + G(\theta^{new}) \right) = 1 - \sum_{t \in \mathcal{T}_g} \pi_{gtc}^{new}.$$

This leads to the following system of equations

$$\pi_{gtc}^{new} = \sum_{i \in \mathcal{C}_c} (\pi_{gtc}^{old} Y_{gi} + \gamma_{gt}^{old} - 1) \frac{1}{\lambda}$$

$$\sum_{t \in \mathcal{T}_g} \pi_{gtc}^{new} = 1.$$

Combining both equations gives

$$\sum_{i \in \mathcal{C}_c} \sum_{t \in \mathcal{T}_g} (\pi_{gtc}^{old} Y_{gi} + \gamma_{gt}^{old} - 1) \frac{1}{\lambda} = 1$$

and so

$$\lambda = \sum_{i \in \mathcal{C}_c} \sum_{t \in \mathcal{T}_g} (\pi_{gtc}^{old} Y_{gi} + \gamma_{gt}^{old} - 1),$$

finally giving our update step for $\pi_{gtc}$,

$$\pi_{gtc}^{new} = \frac{\sum_{i \in \mathcal{C}_c} (\pi_{gtc}^{old} Y_{gi} + \gamma_{gt}^{old} - 1)}{\sum_{t \in \mathcal{T}_g} \sum_{i \in \mathcal{C}_c} (\pi_{gtc}^{old} Y_{gi} + \gamma_{gt}^{old} - 1)}. \tag{20}$$

The expression in Equation (20) strikes an intuitive balance between the observed data and prior distribution. First, if no prior distribution is available, $\gamma_{gt}^{old} = 1$ and the expression reduces to a ratio of the estimated number of molecules produced by TF $t$ over all molecules produced for gene $g$, i.e., the standard Multinomial estimating equation. If a prior distribution is used, either known or estimated, the parameters of the prior can be seen as pseudocounts that inform the estimation of $\pi_{gtc}$.

Once we have updated our estimates for $\pi_{gtc}$, we update our estimates for $\bar{\mu}_{.tc}$ as

$$\bar{\mu}_{.tc}^{new} = \frac{1}{|\mathcal{G}_t|} \sum_{g \in \mathcal{G}_t} \mu_{g.c} \pi_{gtc}^{new}.$$

**Repressions.** If significant repressions between TFs are found, we multiply the estimated $\bar{\mu}_{.tc}^{new}$ by $\rho_{tc}$, where $\rho_{tc}$ is estimated as detailed in Section 2.2.2.

**M-step II: Estimate $\gamma_{gt}^{new}$, given $\pi_{gtc}^{new}$.** There is no closed-form solution for this update step, as the $\gamma_{gt}$ parameters are also involved in the gamma functions in $Q$. However, since we have already estimated $\pi_{gtc}$, we can readily estimate the relative magnitude of the $\gamma_{gt}$ parameters, i.e.,

$$\tilde{\gamma}_{gt.}^{new} = \frac{\gamma_{gt}^{new}}{\sum_{t \in \mathcal{T}_g} \gamma_{gt}^{new}} = \frac{\bar{\pi}_{gt}^{new}}{\sum_{t \in \mathcal{T}_g} \bar{\pi}_{gt}^{new}},$$

where $\bar{\pi}_{gt}^{new} = \sum_c \pi_{gtc}^{new}/C$ and $C$ is the number of cell types. This means that we only need to estimate the precision $\gamma_{g.}^{new} = \sum_{t \in \mathcal{T}_g} \gamma_{gt}^{new}$. We use a closed-form approximate MLE update step as provided in Minka [2012],

$$\gamma_{g.}^{new} = \frac{(|\mathcal{T}_g| - 1)/2}{- \sum_{t \in \mathcal{T}_g} \tilde{\gamma}_{gt.}^{new} \left\{ \frac{1}{C} \sum_{c=1}^{C} \log \left( \frac{\pi_{gtc}^{new}}{\tilde{\gamma}_{gt.}^{new}} \right) \right\}}.$$

**Scaling of $\gamma_{gt}$.** Equation (20) shows that the prior paramaters $\gamma_{gt}$ can be considered as pseudocounts in the update step for $\pi_{gtc}$. It is therefore sensible to consider scaling $\gamma_{gt}$ with respect to the data $\mathbf{Y}$, in order to balance the contribution of the prior vs. the data. Providing equal weight to the data and the prior would correspond to $\sum_{i\in\mathcal{C}_c} Y_{gi} = \sum_t A_{gt}\gamma_{gt}$, i.e., the total observed molecules for a gene $g$ is equal to the magnitude of the Dirichlet parameters. This holds because there is an equal amount of (pseudo)counts to partition among the transcription factors according to the observed data as well as the prior distribution, see Equation (20). Note that the scaling is done per cell type, recycling the same $\gamma_{gt}$ parameters, as estimation happens for each cell type separately. Given a scaling factor $S$, which defines the relative prior vs. data weight ($S = 1$ if equal weight), we can rescale $\gamma_{gt}$ as

$$\gamma_{gtc}^S = \gamma_{gt} S \frac{Y_{g.c}}{\sum_t A_{gt}\gamma_{gt}}.$$

Note that now the prior parameters are cell-type-specific upon scaling. If scaling occurs, the $\gamma_{gt}$ parameters are thus replaced by the appropriate $\gamma_{gtc}^S$ parameters in the update steps of the EM algorithm.

In the software, this scaling is implemented in the `alphaScale` argument for the hierarchical Poisson model, where the prior distribution is assumed known. By default, $S = 1$, i.e., equal prior and data weight. For $S > 1$, the prior has more weight than the observed data, and vice versa for $S < 1$.

**Stopping rule.** Stop when the log-likelihood increments are below a user-defined threshold (set at 0.01 by default). Other stopping rules, like tracking convergence of parameter estimates using a norm of their differences, could be used but are not implemented. The user can investigate convergence of the EM algorithm by plotting the trace of the log-likelihood as a function of EM iterations. If the EM algorithm does not converge, it will stop after a user-specified maximum number of iterations.

**Implementation.** While the EM algorithm here is written in terms of the full data $Y_{gi}$, in practice it is implemented using the sufficient statistics for the Poisson distribution $Y_{gc} = \sum_{i\in\mathcal{C}_c} Y_{gi}$, leading to considerable computational gains in terms of memory consumption and speed.

**Interpretation.** For downstream interpretation and differential activity testing, we can work with the expected number of molecules produced by TF $t$ in cell $i$ from cell type $c$, i.e.,

$$E\left[\sum_{g\in\mathcal{G}_t} Z_{gti} \middle| Y_{gi}\right] = \sum_{g\in\mathcal{G}_t} E[Z_{gti}|Y_{gi}] = \sum_{g\in\mathcal{G}_t} Y_{gi}\pi_{gtc(i)}. \tag{21}$$

### 6 Model identifiability and examples using small networks

Using only the basic model assumptions (A1) and (A2), we explore the identifiability of our statistical model, where we consider the model identifiable if existence and uniqueness of its parameters holds, that is, there cannot be a different set of $\alpha_{gc}$ and $\beta_{tc}$ parameter values that can yield the same distribution for the observed data $Y_{gi}$, i.e., the same $\mu_{gtc}$.

Given assumptions (A1) and (A2), the model is identifiable if and only if the following set of equations holds and has a unique solution for each cell type $c$

$$\begin{aligned} \alpha_{gc}\sum_{t=1}^{T}\beta_{tc}A_{gt} &= \mu_{g.c}, \quad \forall g \\ \frac{\sum_g \alpha_{gc}A_{gt}}{\sum_g A_{gt}} &= 1, \quad \forall t. \end{aligned} \tag{22}$$

This corresponds to a set of $G + T$ equations for the $G + T$ $\alpha$ and $\beta$ parameters, where $G$ equations are quadratic and $T$ are linear. Note that these equations are the method-of-moments estimating equations.

While the system in Equation (22) seems simple, there are no obvious conditions under which the solution exists and is unique. The GRN $\mathbf{A}$ is, however, crucial for the identifiability of the model. Indeed, we can use the structure of the GRN to determine sufficient conditions for identifiability. In the trivial case where each gene is regulated by a single TF and each TF regulates a single gene (i.e., $\mathbf{A}$ can be rewritten as the identity matrix by reordering its rows or columns), we have $\alpha_{gc} = 1 \ \forall g$ and $\beta_{tc} = \mu_{g(t).c} \ \forall t$, where $g(t)$ is defined as the unique gene such that $A_{g(t)t} = 1$, and thus an identifiable model.

We posit necessary, but not sufficient, identifiability conditions in Methods Section 2.1.4. Indeed, if one condition does not hold, the system of equations automatically becomes underdetermined. Furthermore, the examples below also demonstrate the restrictions imposed by the model assumptions, where in more complex settings the (ratios of) TF activities are constrained according to (ratios of) gene expression; in particular, see Example 4 below.

The core set of sufficient conditions for identifiability for a general GRN $\mathbf{A}$ remains an open problem. Indeed, adhering to the above two minimal conditions described in Section 2.1.4 does not guarantee identifiability.

We can use small networks as examples to gain intuition on identifiability conditions under which the system of equations in (22) has a unique solution. Below, we establish model identifiability for three small network structures. However, we also show that another small network structure leads to unidentifiable parameters. Without loss of generality, we will be assuming that all cells are from a single cell population and will henceforth drop the subscript $c$.

#### Example 1: One TF, one gene

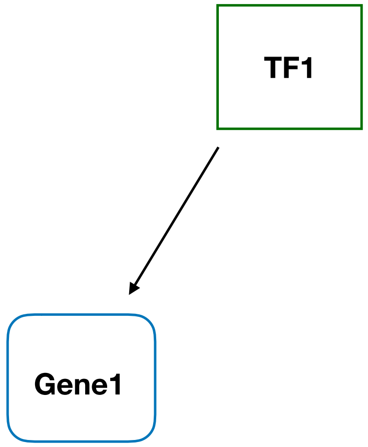


Supplementary Methods Figure 1: *One TF, one gene.*

In the example shown in Supplementary Methods (SM) Figure 1, $G = 1$ and $T = 1$; the smallest GRN possible. The adjacency matrix is a scalar $(1 \times 1)$, $\mathbf{A} = (1)$. The system of equations as derived from Equation (22) is

$$\begin{aligned} \mu_1 &= \alpha_1 \beta_1 \\ \alpha_1 &= 1. \end{aligned}$$

This straightforwardly leads to $\beta_1 = \mu_1$ (note that we assume $\mu_1$ is known or can be inferred from the observed data $\mathbf{Y}$ as in Equation (14)). This is a sensible solution: Since TF1 only regulates a single gene, Gene1, and that gene is uniquely regulated by TF1, its activity is equal to the gene expression of Gene1. If the gene is highly expressed, the TF will be highly active and vice versa.

#### Example 2: One TF, two genes

In the example shown in SM Figure 2, $G = 2$ and $T = 1$. The $2 \times 1$ adjacency matrix is

$$\mathbf{A} = \begin{pmatrix} 1 \\ 1 \end{pmatrix}.$$

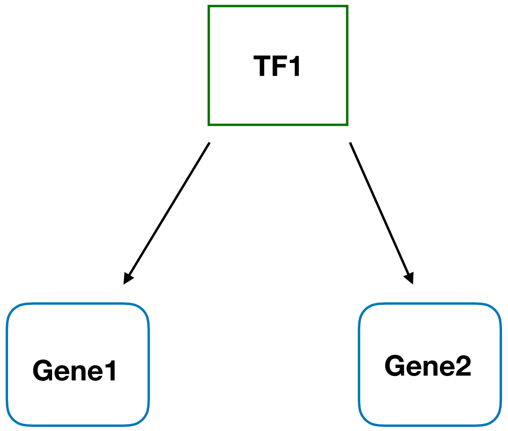


Supplementary Methods Figure 2: *One TF, two genes.*

The system from Equation (22) is

$$\begin{aligned}\mu_1 &= \alpha_1\beta_1\\ \mu_2 &= \alpha_2\beta_1\\ \alpha_1+\alpha_2 &= 2\end{aligned}$$

and can be rewritten as

$$\begin{aligned}\mu_1 &= 2\beta_1-\alpha_2\beta_1\\ \alpha_2 &= \mu_2/\beta_1\\ \alpha_1 &= 2-\alpha_2,\end{aligned}$$

which yields solutions

$$\begin{aligned}\beta_1 &= \frac{\mu_1+\mu_2}{2}\\ \alpha_1 &= \frac{2\mu_1}{\mu_1+\mu_2}\\ \alpha_2 &= \frac{2\mu_2}{\mu_1+\mu_2}.\end{aligned}$$

To interpret this result, we can rewrite $\alpha_1+\alpha_2$ using (A1) as $\frac{\mu_1}{\beta_1}+\frac{\mu_2}{\beta_1}=2$ and so $\mu_1+\mu_2=2\beta_1$. This is again a sensible result, as we can interpret $\beta_1$ as the average number of molecules produced by TF1 across all the genes it is regulating, given (A2).

#### Example 3: Two TFs, one gene

In the example shown in SM Figure 3, $G=1$ and $T=2$. This is an example where $T>G$. The $1\times 2$ adjacency matrix is

$$\mathbf{A}=\begin{pmatrix}1 & 1\end{pmatrix}.$$

The system from Equation (22) is

$$\begin{aligned}\mu_1 &= \alpha_1(\beta_1+\beta_2)\\ \alpha_1 &= 1.\end{aligned}$$

There are only two independent equations and three unknowns. This system is underdetermined and there is no unique solution, therefore rendering the model unidentifiable. In terms of our application, this makes sense since we have no grounds by which we may differentially partition the molecules from Gene1 to TF1 and TF2.

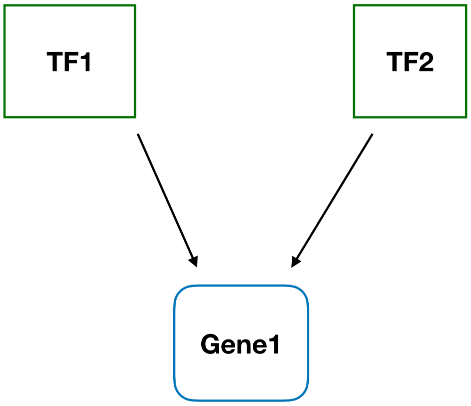


Supplementary Methods Figure 3: *Two TFs, one gene.*

##### Example 4: Two TFs, three genes

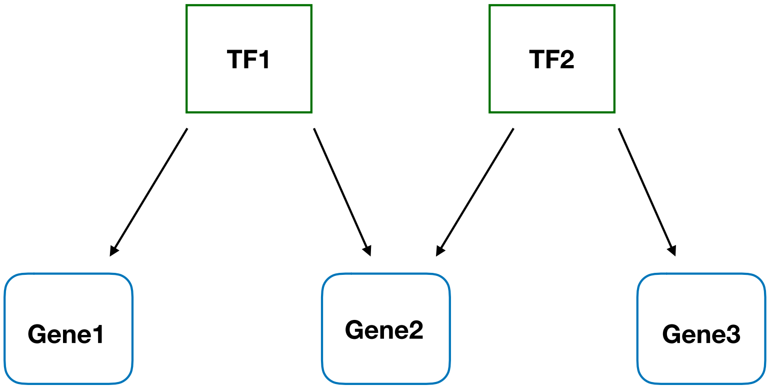


Supplementary Methods Figure 4: *Two TFs, three genes.*

In the example shown in SM Figure 4, $G = 3$ and $T = 2$. The $3 \times 2$ adjacency matrix is

$$\mathbf{A} = \begin{pmatrix} 1 & 0 \\ 1 & 1 \\ 0 & 1 \end{pmatrix}.$$

The system from Equation (22) is

$$\begin{aligned} \mu_1 &= \alpha_1\beta_1 \\ \mu_2 &= \alpha_2(\beta_1 + \beta_2) \\ \mu_3 &= \alpha_3\beta_2 \\ 2 &= \alpha_1 + \alpha_2 \\ 2 &= \alpha_2 + \alpha_3 \end{aligned}$$

and can be rewritten as

$$\begin{aligned}
\beta_1 &= \frac{\mu_1}{2-\alpha_2} \\
\mu_2 &= \alpha_2 \left( \frac{\mu_1}{2-\alpha_2} + \frac{\mu_3}{2-\alpha_2} \right) \\
\beta_2 &= \frac{\mu_3}{2-\alpha_2} \\
\alpha_1 &= 2-\alpha_2 \\
\alpha_3 &= 2-\alpha_2.
\end{aligned}$$

The second equation can be solved as $\alpha_2 = \frac{2\mu_2}{\mu_1+\mu_2+\mu_3}$ and, since all other equations are a function of $\alpha_2$, the system is solved.

To interpret this result, note that the equations

$$\begin{aligned}
2 &= \alpha_1 + \alpha_2 \\
2 &= \alpha_2 + \alpha_3
\end{aligned}$$

imply $\alpha_1 = \alpha_3$. Using (A1), this yields $\frac{\mu_1}{\beta_1} = \frac{\mu_3}{\beta_2}$ and so we have the constraint that $\frac{\mu_1}{\mu_3} = \frac{\beta_1}{\beta_2}$. This is a sensible constraint. Indeed, the ratio of TF activities between TF1 and TF2 is reflected by the respective expression of Gene1 and Gene3, as these genes are uniquely regulated by these TFs. This will then also drive the partitioning of molecules of Gene2 according to Equation (9).

### 7 Supplementary Figures

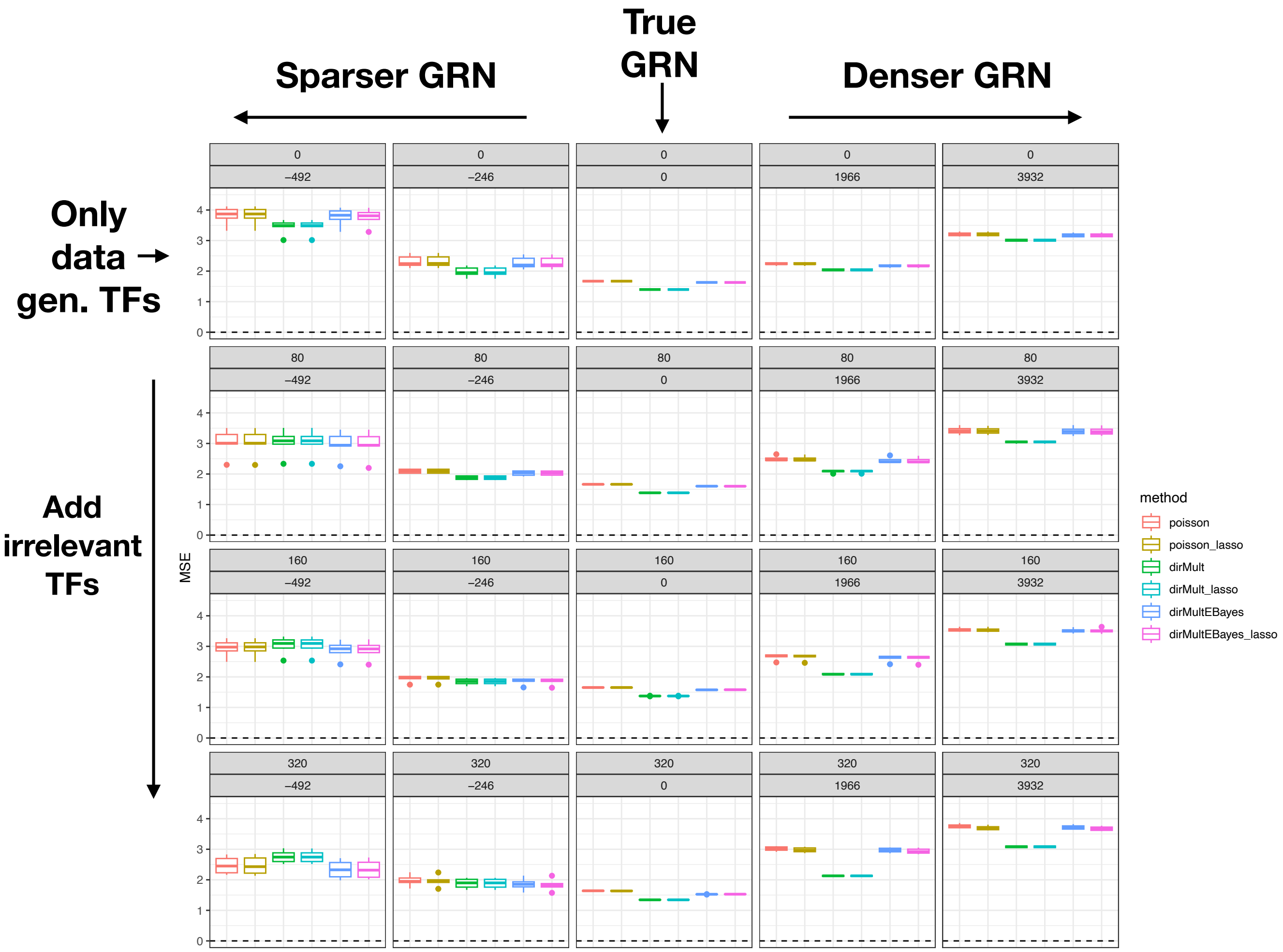


Supplementary Figure 1: *Gamma-model-based simulation study results: Estimation accuracy.* Six methods were evaluated on each of six simulated datasets. For each method, the mean squared error (MSE) for estimating the TF activities $m\bar{u}_{.tc}$ was computed for each of six datasets and the six MSE values were summarized using a boxplot. The different panels represent different GRNs used as input to each method. The first row corresponds to a GRN where only TFs that were involved in the data-generating process are used. Rows 2–4 are rows where, respectively, 80, 160, and 320 'irrelevant' TFs (i.e., TFs that did not contribute to the data-generating process and regulate a random set of genes) are added to the GRN. The different columns represent removing or adding edges to the true GRN. In the middle column, we use the true GRN. Moving to the left (right) of the middle column, we remove (add) true edges to the GRN. Panel subtitles indicate the number of irrelevant TFs added to the GRN and edges added/removed from the GRN.

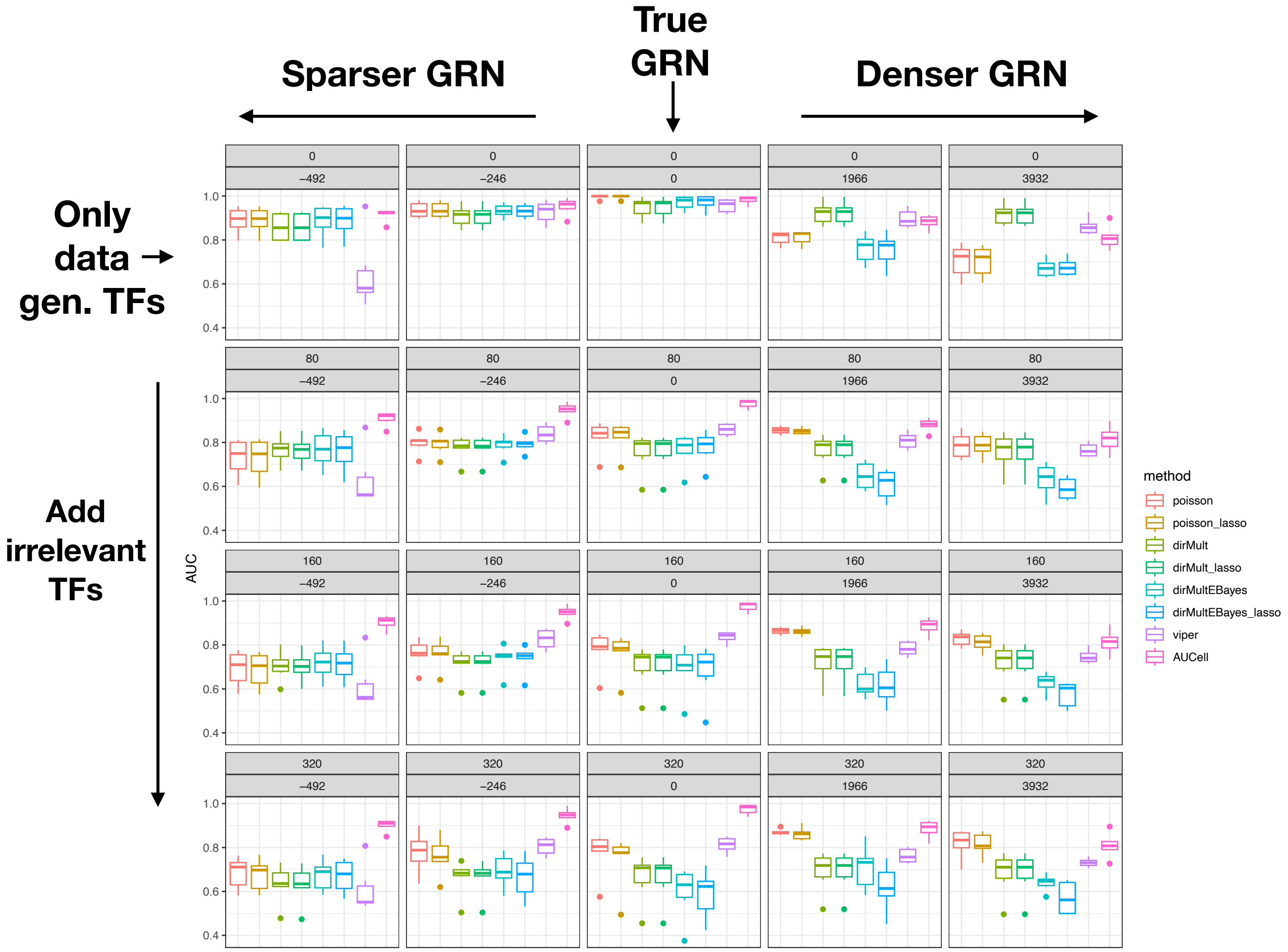


Supplementary Figure 2: *Gamma-model-based simulation study results: Performance when using limma(-voom) for downstream analysis.* Eight methods were evaluated on each of six simulated datasets. For each method and dataset, the area under the receiver operating characteristic curve (AUC) for detecting the TFs that are differentially active between the cell types was computed; either limma or limma-voom was used for the differential activity analysis. For each method, the six AUCs were summarized using a boxplot. The different panels represent different GRNs used as input to each method. The first row corresponds to a GRN where only TFs that were involved in the data-generating process are used. Rows 2–4 are rows where, respectively, 80, 160, and 320 'irrelevant' TFs (i.e., TFs that did not contribute to the data-generating process and regulate a random set of genes) are added to the GRN. The different columns represent removing or adding edges to the true GRN. In the middle column, we use the true GRN. Moving to the left (right) of the middle column, we remove (add) true edges to the GRN. Panel subtitles indicate the number of irrelevant TFs added to the GRN and edges added/removed from the GRN.

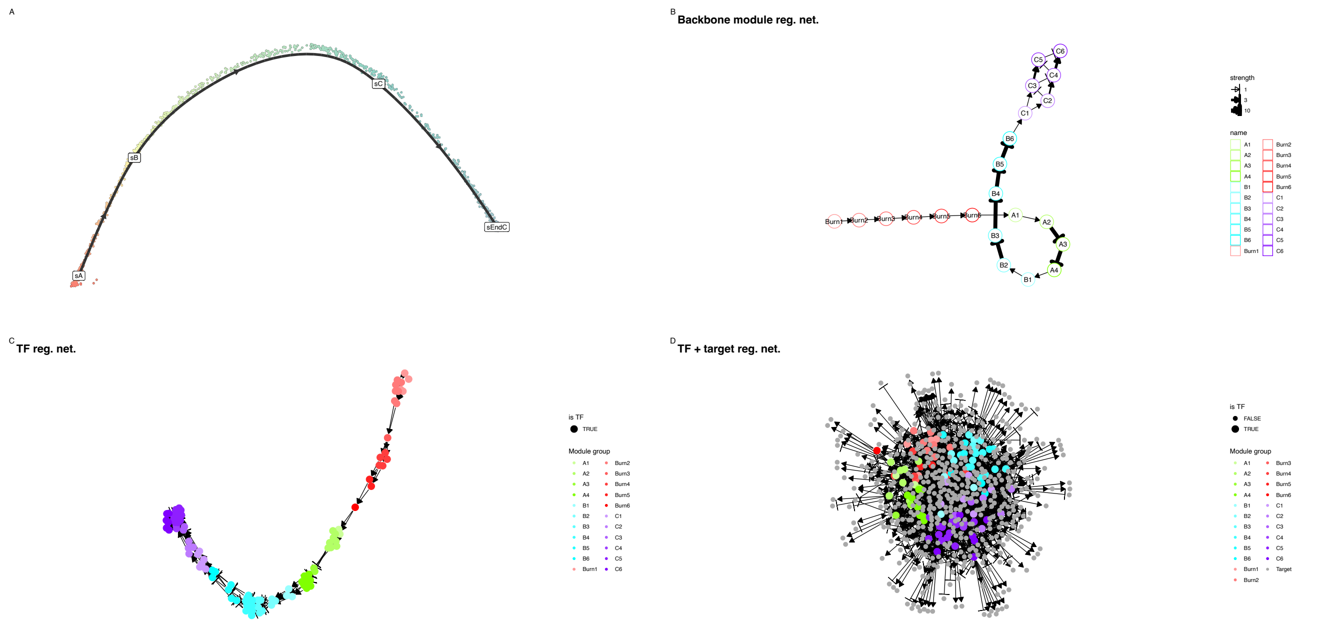


Supplementary Figure 3: *Example gene regulatory network for the* `dyngen` *simulated datasets.* Each node in the network represents a module, transcription factor or gene. A module is a set of genes that can regulate each other, and a module can also regulate other modules. Edges between nodes correspond to interactions between the nodes. An arrow represents a positive interaction (activation), while a blunt arrow represents a negative interaction (repression). **(a)** Simulated trajectory in reduced-dimensional space using multidimensional scaling. Each point represents a single cell, and the black line represents the trajectory, with the different modules indicated by boxes. **(b)** The interactions between the modules that are used to construct the GRNs. Each module consists of a few transcription factors, and is used as a backbone to construct the GRN. **(c)** The transcription factor regulatory network. **(d)** The full gene regulatory network.

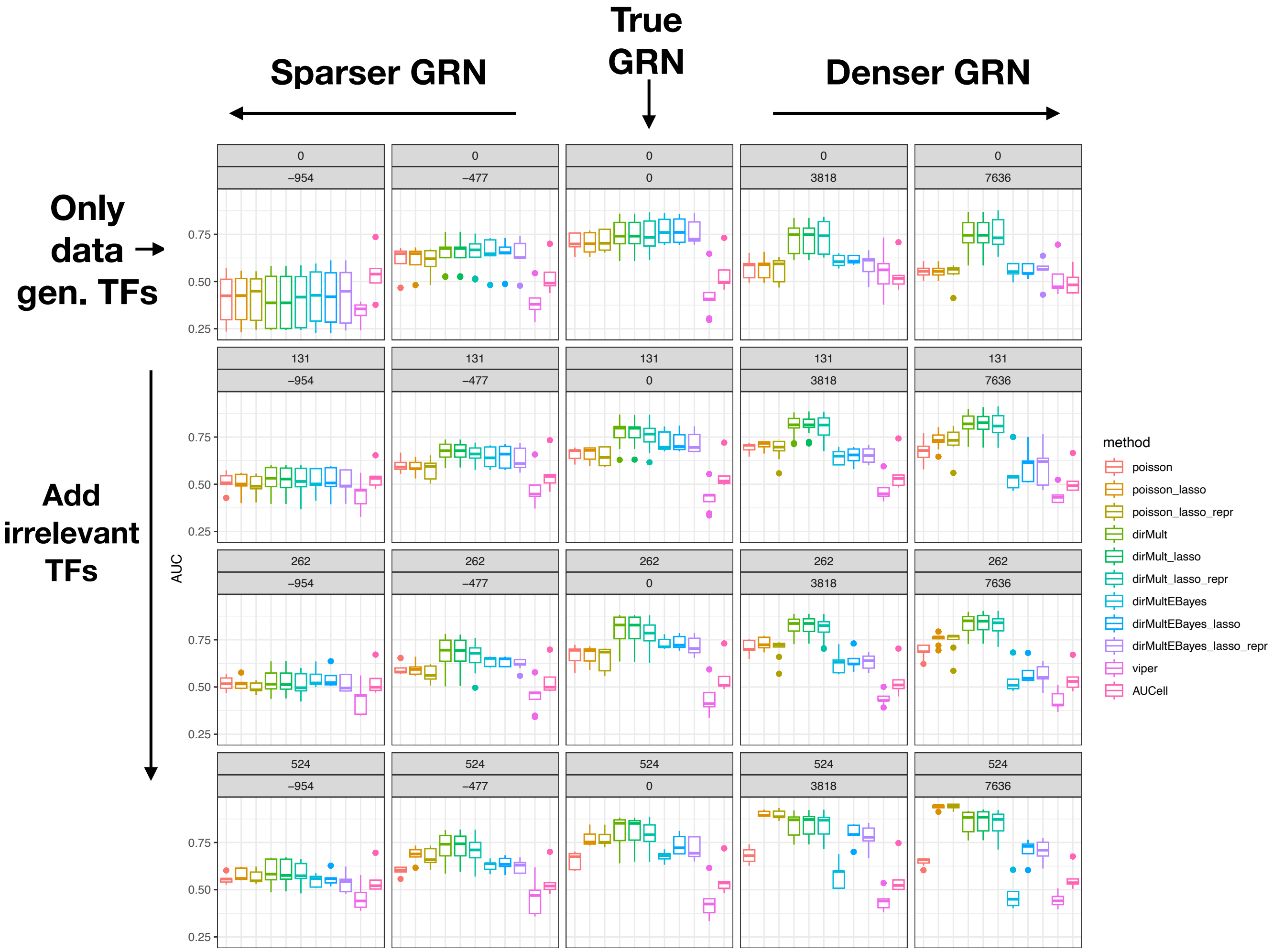


Supplementary Figure 4: *`dyngen` simulation study results, using a prior that discriminates better between true and false positive edges.* Eleven methods were evaluated on each of six simulated datasets. For each method and dataset, the area under the receiver operating characteristic curve (AUC) for detecting the TFs that are differentially active between the cell types was computed. For each method, the six AUCs were summarized using a boxplot. The different panels represent different GRNs used as input to each method. The first row corresponds to a GRN where only TFs that were involved in the data-generating process are used. For rows 2–4, respectively, $131, 262$, and 524 'irrelevant' TFs (i.e., TFs that do not contribute to the data-generating process and regulate a random set of genes) were added to the GRN. The different columns represent removing or adding edges to the true GRN. In the middle column, we use the true GRN. Moving to the left (right) of the middle column, we remove (add) edges to the true GRN. Panel subtitles indicate the number of irrelevant TFs added to the GRN and edges added/removed from the GRN.

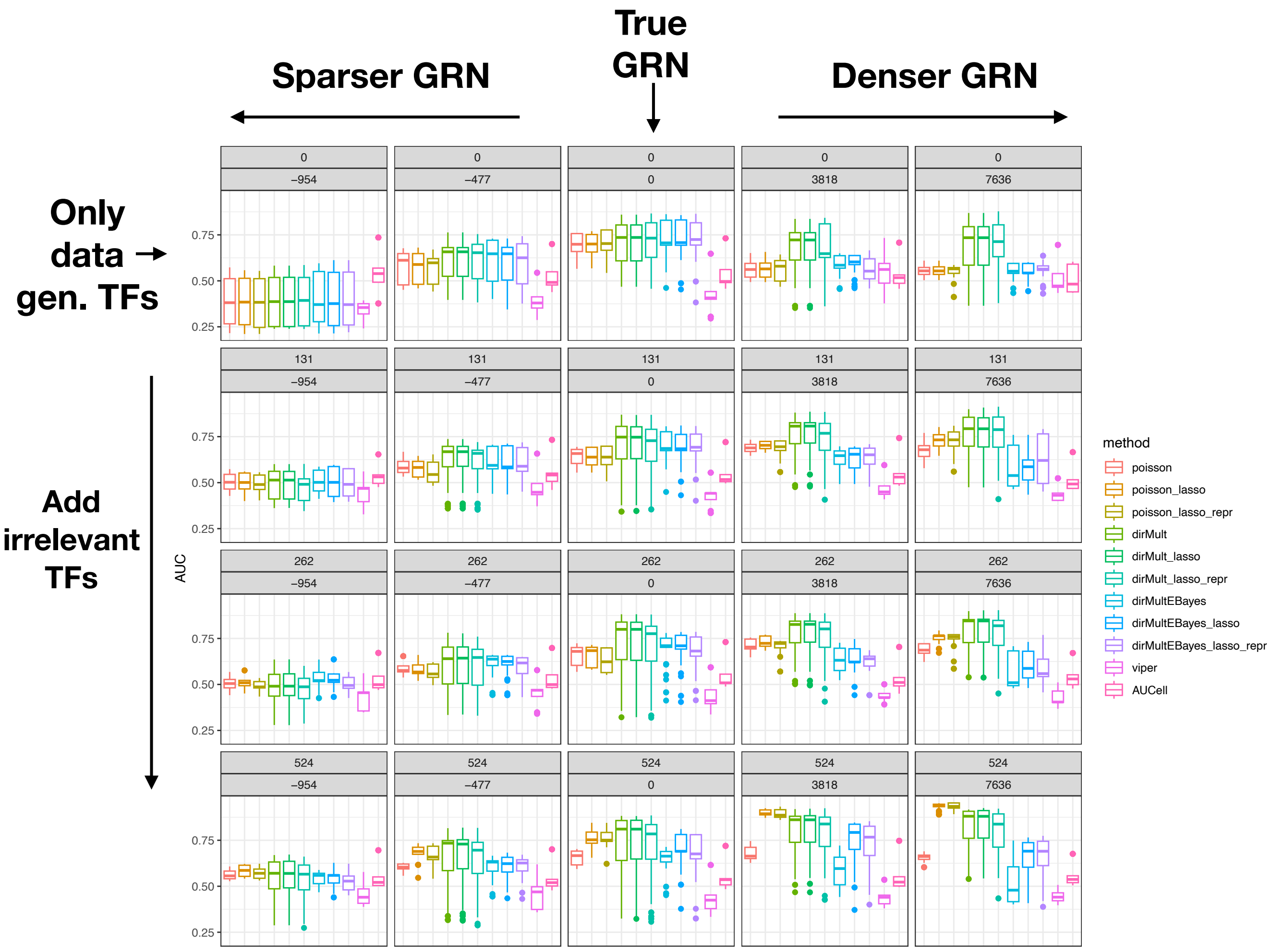


Supplementary Figure 5: *`dyngen` simulation study results, using limma(-voom) for differential activity analysis.* Eleven methods were evaluated on each of six simulated datasets. For each method and dataset, the area under the receiver operating characteristic curve (AUC) for detecting the TFs that are differentially active between the cell types was computed. For each method, the six AUCs were summarized using a boxplot. The different panels represent different GRNs used as input to each method. The first row corresponds to a GRN where only TFs that were involved in the data-generating process are used. For rows 2–4, respectively, 131, 262, and 524 'irrelevant' TFs (i.e., TFs that do not contribute to the data-generating process and regulate a random set of genes) were added to the GRN. The different columns represent removing or adding edges to the true GRN. In the middle column, we use the true GRN. Moving to the left (right) of the middle column, we remove (add) edges to the true GRN. Panel subtitles indicate the number of irrelevant TFs added to the GRN and edges added/removed from the GRN.

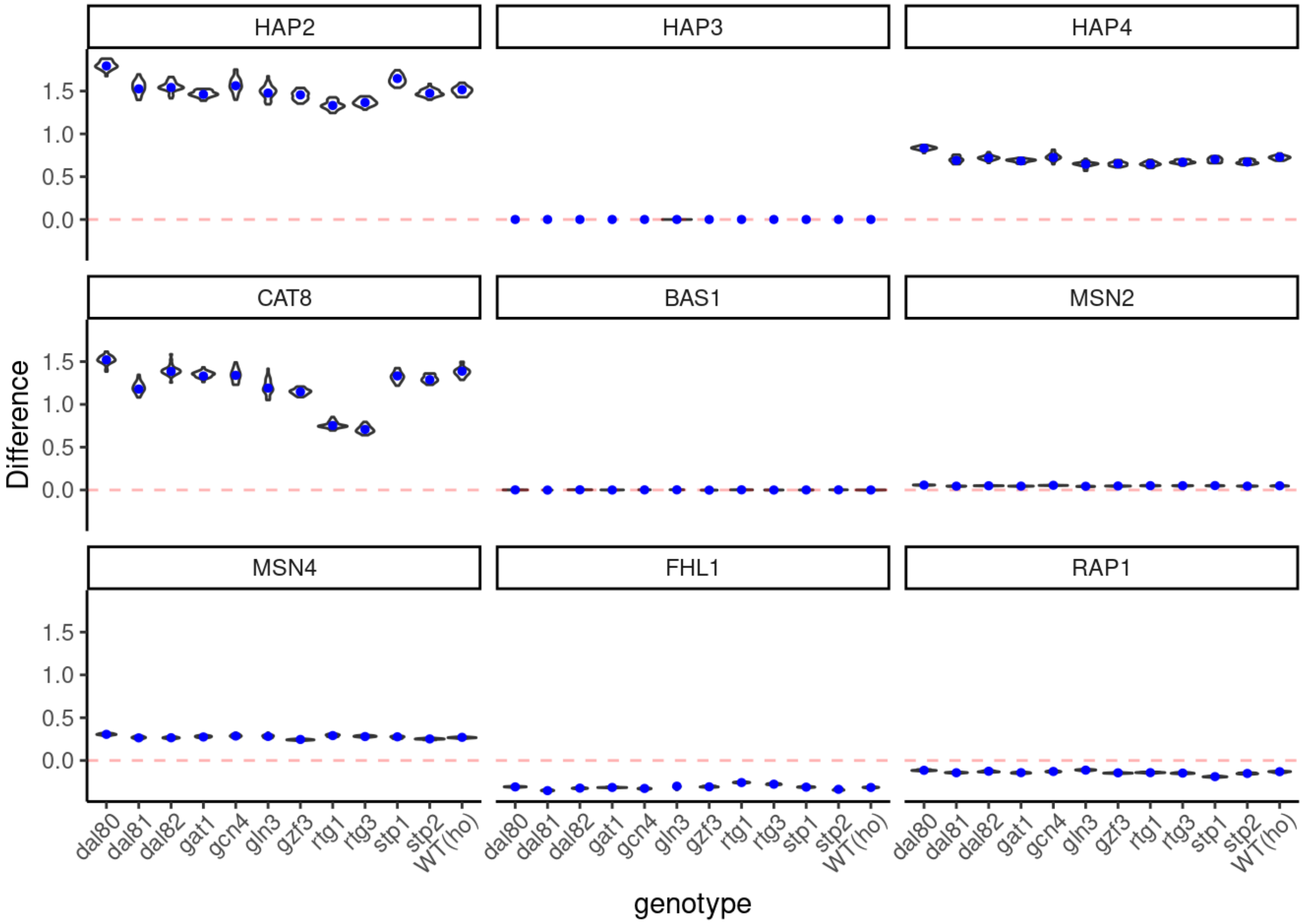


Supplementary Figure 6: *Yeast knock-out case study: TF activity differences for nine TFs known to be involved in the diauxic shift, across cell bootstraps.* Each panel corresponds to a transcription factor, and the difference between TF activities in the diauxic shift vs. YPD conditions are shown (y-axis) for the different genotypes (x-axis). Each difference is calculated as $\hat{\bar{\mu}}_{.t,diauxic} - \hat{\bar{\mu}}_{.t,YPD}$, with $\hat{\bar{\mu}}_{.t,diauxic}$ the estimated activity for TF $t$ in the diauxic condition, and similar for the YPD condition. The blue dot denotes the estimated difference on the full data, while the violin plots show the variability across 30 cell bootstraps.

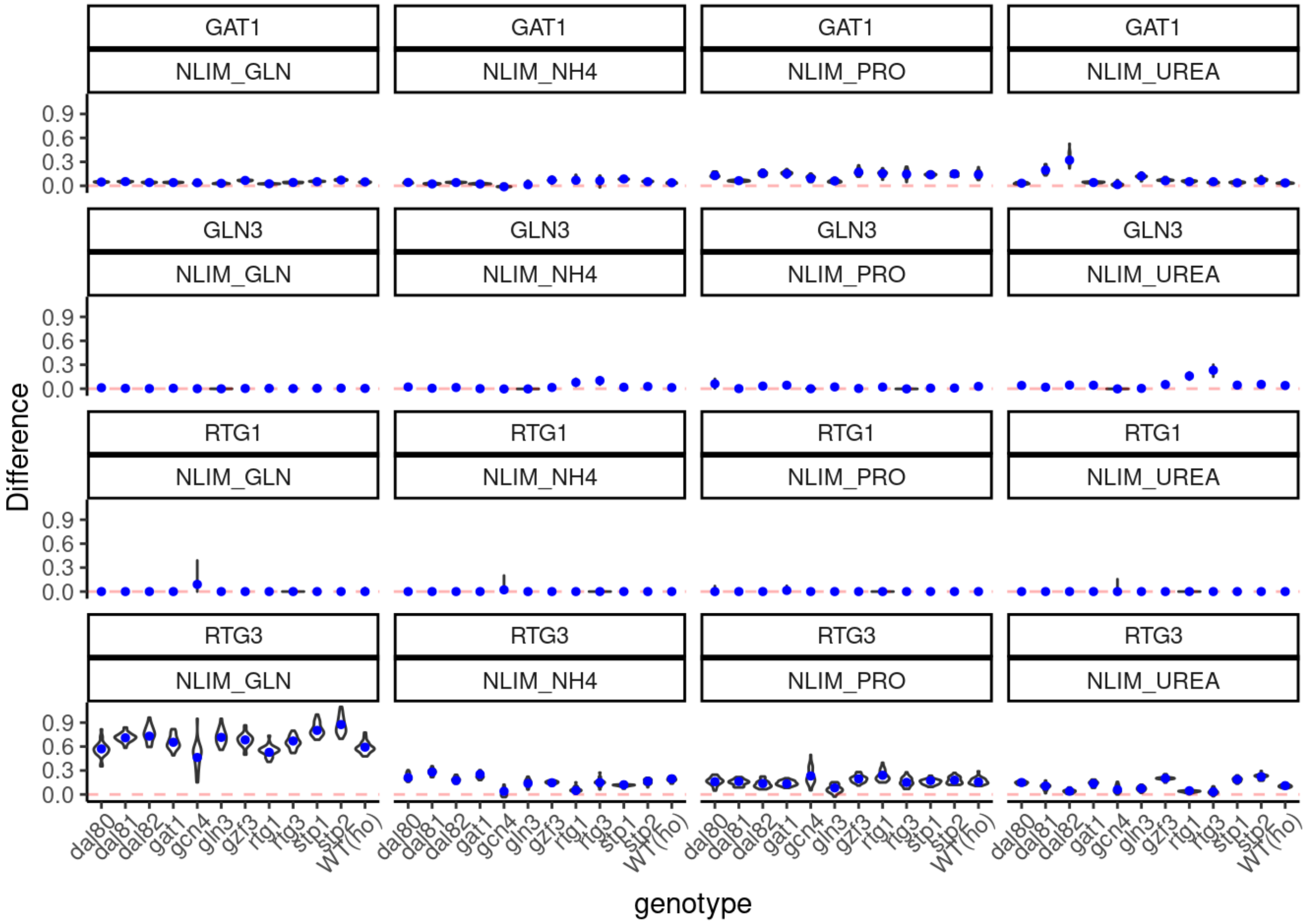


Supplementary Figure 7: *Yeast knock-out case study: TF activity differences for four TFs known to be involved in nitrogen metabolosim, across cell bootstraps.* Each panel corresponds to a transcription factor in a particular nitrogen limiting condition, and the differences between TF activities in that respective condition vs. the YPD condition are shown (y-axis) for the different genotypes (x-axis). Each difference is calculated as $\hat{\bar{\mu}}_{.t,NLIM} - \hat{\bar{\mu}}_{.t,YPD}$, with $\hat{\bar{\mu}}_{.t,NLIM}$ the estimated activity for TF $t$ in one of the nitrogen limiting conditions, and similar for the YPD condition. The blue dot denotes the estimated difference on the full data, while the violin plots show the variability across 30 cell bootstraps.

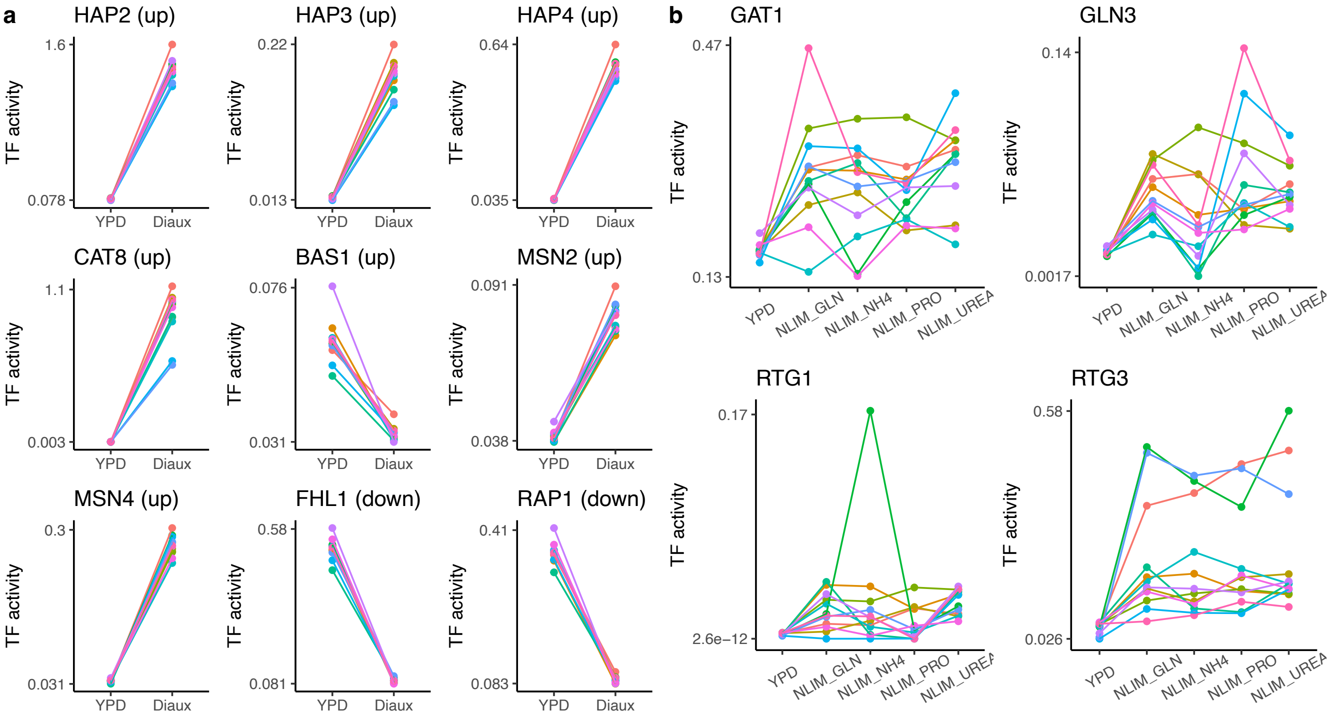


Supplementary Figure 8: *Yeast knock-out case study: TF activities, estimated using the hierarchical Poisson model.* **(a)** TF activity estimates for nine TFs involved in the diauxic shift. Each color represents a different gene deletion mutant. **(b)** TF activity estimates for four TFs involved in nitrogen metabolism. Each color represents a different gene deletion mutant.

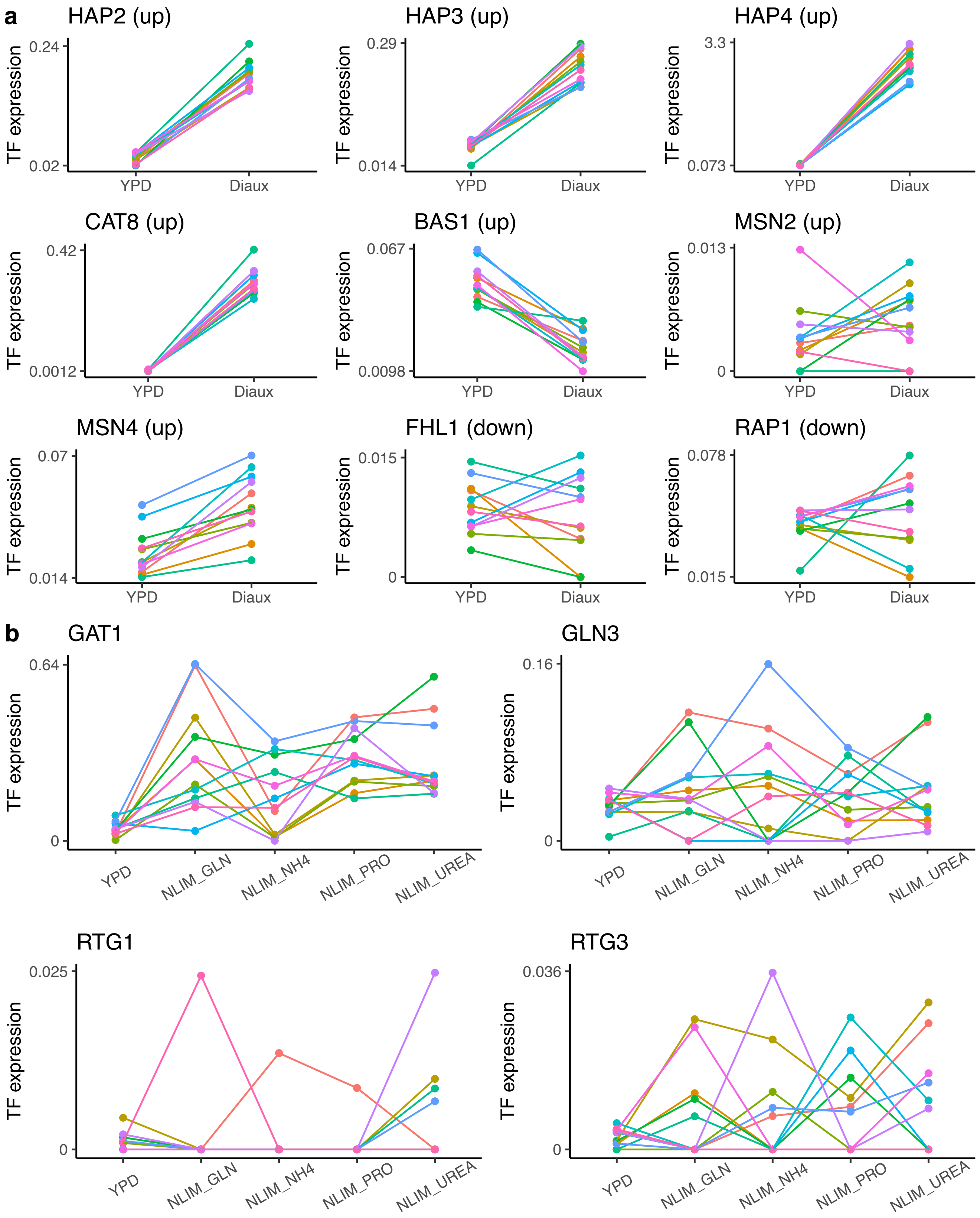


Supplementary Figure 9: *Yeast knock-out case study: TF expression.* **(a)** TF expression averages for nine TFs involved in the diauxic shift. **(b)** TF expression averages for four TFs involved in nitrogen metabolism. Each color represents a different gene deletion mutant.

### 8 Supplementary Tables

Supplementary Table 1: Olfactory epithelium case study: Gene set enrichment results for the genes regulated by transcription factors found to be most active in the activated HBC stage. The top 20 gene sets are shown, and results were obtained using `gProfiler` [Raudvere et al., 2019].

| | Gene set |
|---|---|
| 1 | developmental process |
| 2 | anatomical structure development |
| 3 | system development |
| 4 | localization |
| 5 | positive regulation of biological process |
| 6 | positive regulation of cellular process |
| 7 | tissue development |
| 8 | anatomical structure morphogenesis |
| 9 | cellular response to chemical stimulus |
| 10 | multicellular organism development |
| 11 | animal organ development |
| 12 | response to organic substance |
| 13 | negative regulation of cellular process |
| 14 | negative regulation of biological process |
| 15 | cellular response to organic substance |
| 16 | cell death |
| 17 | regulation of signal transduction |
| 18 | organonitrogen compound metabolic process |
| 19 | cell migration |
| 20 | cellular process |

Supplementary Table 2: Olfactory epithelium case study: Gene set enrichment results for the genes regulated by transcription factors found to be most active in the GBC stage. The top 20 gene sets are shown, and results were obtained using `gProfiler` [Raudvere et al., 2019].

| | Term name |
|---|---|
| 1 | cellular metabolic process |
| 2 | cellular nitrogen compound metabolic process |
| 3 | DNA replication |
| 4 | cell cycle |
| 5 | macromolecule metabolic process |
| 6 | organic substance metabolic process |
| 7 | nitrogen compound metabolic process |
| 8 | heterocycle metabolic process |
| 9 | cellular aromatic compound metabolic process |
| 10 | nucleobase-containing compound metabolic process |
| 11 | organic cyclic compound metabolic process |
| 12 | metabolic process |
| 13 | cell cycle process |
| 14 | primary metabolic process |
| 15 | chromosome organization |
| 16 | DNA metabolic process |
| 17 | cellular macromolecule metabolic process |
| 18 | organelle organization |
| 19 | mitotic cell cycle |
| 20 | nucleic acid metabolic process |

Supplementary Table 3: Olfactory epithelium case study: Gene set enrichment results for the genes regulated by transcription factors found to be most active in the neuronal stage. The top 20 gene sets are shown, and results were obtained using `gProfiler` [Raudvere et al., 2019].

| | Term name |
|---|---|
| 1 | cellular component organization |
| 2 | cellular component organization or biogenesis |
| 3 | cellular metabolic process |
| 4 | cellular localization |
| 5 | protein localization |
| 6 | plasma membrane bounded cell projection organization |
| 7 | cell projection organization |
| 8 | organelle organization |
| 9 | cellular macromolecule localization |
| 10 | cellular protein localization |
| 11 | cellular component assembly |
| 12 | macromolecule localization |
| 13 | cellular component biogenesis |
| 14 | establishment of localization in cell |
| 15 | cellular macromolecule metabolic process |
| 16 | establishment of protein localization |
| 17 | cellular process |
| 18 | intracellular transport |
| 19 | protein transport |
| 20 | nervous system development |
